# IRF2 degradation tunes the innate immune response

**DOI:** 10.1101/2025.09.23.678148

**Authors:** Cristhian Cadena, Rohit Reja, Emma Bolech, Joshua D. Webster, Vasumathi Kameswaran, Marco de Simone, Cynthia Chen, Jian Jiang, Kathy Hotzel, Kamela Alegre, Zhenyu Tan, Raymond Newland, Ryan Kelly, Spyros Darmanis, Bence Daniel, Ishan Deshpande, Kim Newton, Nobuhiko Kayagaki, Vishva M. Dixit

## Abstract

The transcription factor IRF2 protects against skin inflammation in mice and humans but, paradoxically, promotes pyroptosis by inducing *Gsdmd*. How IRF2 activates some proinflammatory genes, but suppresses others is unclear. We show that skin inflammation in *Irf2*-deficient mice is driven by IRF1 activation of interferon-stimulated genes (ISGs). Chromatin profiling revealed that IRF1 and IRF2 occupy the same ISG regulatory sites, but as a weaker transcriptional activator, IRF2 limited ISG transcription by IRF1. Toll-like receptor signaling favored IRF1-driven transcription by inducing *Irf1*. In addition, IRF1 recruited the ubiquitin ligase SPOP to ISG sites, resulting in proteasomal degradation of IRF2. This shift from IRF2 to IRF1 occupancy enhanced ISG transcription. Collectively, these findings define a hierarchical transcriptional circuit in which IRF2 limits IRF1 activity under homeostatic conditions but is displaced during an immune response, allowing IRF1-dependent gene programs central to innate immunity and autoinflammation.

## Introduction

Interferons (IFNs) and ISGs mediate innate immune responses to invading pathogens ^1^, but their dysregulation causes autoimmunity ^2^. The interferon regulatory family (IRF) of transcription factors has 9 members with diverse roles in the induction of proinflammatory genes. In the classical model of IFN signaling, sensing of pathogen-associated molecular patterns (PAMPs) by innate immune receptors, including TLRs, triggers signaling cascades that activate IRF3, resulting in transcription of *IFNB1* plus other pro-inflammatory genes ^1^. IRF7, which is closely related to IRF3, also contributes to the expression of IFN-β and IFN-α subtypes (collectively referred to as type I IFNs) in a context and cell-type dependent manner ^3,4^. Type I IFNs secreted by infected cells engage the interferon α/□ receptor (IFNAR) in a paracrine and autocrine manner to activate the JAK-STAT signaling cascade ^1^. Phosphorylated STAT1 and STAT2 interact with IRF9 to form the ISGF3 complex that promotes the expression of hundreds of ISGs, thereby priming cells for antiviral responses ^1^.

While IRF3 and IRF9 are important drivers of IFN-α/□ responses downstream of PRRs, less is known about IRF1 and IRF2. Initial studies suggested that IRF1 activates and IRF2 suppresses *IFNB1* transcription ^5,6,7^, but a later study implicated IRF1 in IFN-γ-induced ISG expression during mycobacterial infections and found it was dispensable for anti-viral signalling ^8^. More recently, IRF1 and the ISGF3 complex were shown to control the opening of IFN-stimulated response element (ISRE)-containing enhancers in response to bacterial lipopolysaccharide (LPS) ^9^. By contrast, IRF2 is still considered an important suppressor of inflammation because its loss causes severe skin inflammation in mice ^10^ and humans ^11,12^.

IRF1 and IRF2 share the same DNA binding motif ^13,14^ and there is overlap in the genes they regulate. Structural studies of the IRF1 and IRF2 DNA binding domains show that both engage the core IRF AAA_NN_GAAA_NNN_AAA motif in a similar manner (where N is any nucleotide) ^13,14^. Both transcription factors minimally bind to the major groove of the GAAA core and the minor groove of the 5′-flanking AA sequence ^13,14^. Given the limited capacity of IRF2 to transcribe *IFNB1* compared to IRF1, it was proposed that IRF1 and IRF2 mutually regulate each other through competitive binding at shared DNA sequences ^5,6,7^. However, the extent to which IRF1 and IRF2 function antagonistically at other genomic sites remains unclear. IRF1 and IRF2 appear to collaborate in the regulation of other target genes. For example, IRF1 and IRF2 both drive the expression of *IL7* and *Tlr3* ^15,16,17^. *Gsdmd* may also be regulated by IRF1 and IRF2 since its expression is reduced but not eliminated by IRF2 deficiency ^18^. The mechanistic details of how cells overcome the suppressive effects of IRF2 when ISG transcription is warranted are unresolved.

Here, we report a role for IRF1 in controlling ISG expression downstream of several IFN-stimulating pathways. We show that transcriptional upregulation of IRF1 promotes IRF2 proteasomal degradation because IRF1 positions the E3 ubiquin ligase SPOP at regulatory sites occupied by IRF2. Eliminating IRF2 enhances IRF1-dependent ISG expression, but also prevents IRF2 from aberrantly activating other proinflammatory genes. This regulatory circuit ensures that potentially deleterious ISGs are expressed only as needed.

## Results

### IRF1 drives GSDMD expression in the absence of IRF2

*Irf2* deficiency suppresses pyroptosis in response to cytoplasmic LPS because cells express less *Gsdmd* than normal ^18^. However, bone marrow-derived macrophages (BMDMs) from *Irf1^−/−^Irf2^−/−^* mice expressed even less *Gsdmd* mRNA and GSDMD protein than *Irf2*^-/-^ BMDMs (Figures S1A and S1B). Similarly, CRISPR/Cas9 knockout of *IRF1* and *IRF2* in primary human macrophages reduced *GSDMD* expression more than knockout of *IRF2* alone (Figures S1C and S1D). Therefore, IRF1 contributes to GSDMD expression in the absence of IRF2.

Chromatin immunoprecipitation-sequencing (ChIP-seq) of wild-type (WT) and *Irf2^-/-^* BMDMs confirmed that IRF2 binds to the *Gsdmd* promoter, where it facilitates acetylation of lysine 27 on histone H3 (H3K27ac), a hallmark of active promoters (Figure S1E). IRF2 is reported to drive expression of *CASP4*, the human orthologue of mouse *Casp11*, in monocytes and macrophages^19^. We detected IRF2 binding at the *Casp11* promoter. However, *Irf2^-/-^* BMDMs maintained H3K27ac at the *Casp11* promoter (Figure S1E) and expressed more *Casp11* than their WT counterparts (Figures S1A and S1B), indicating that IRF2 suppresses *Casp11*. Transcription of *Casp11* was not elevated in *Irf1*^-/-^*Irf2^-/-^*BMDMs (Figures S1A and S1B), suggesting that IRF1 drives *Casp11* expression in the absence of IRF2.

To test whether the IRF2 binding site in the *Gsdmd* promoter ^18^ is the only IRF site controlling *Gsdmd* transcription, we engineered *Gsdmd^Irf/Irf^* mice with nucleotide substitutions at this site (Figure S1F). *Gsdmd^Irf/Irf^* BMDMs resembled *Irf1*^-/-^*Irf2^-/-^*BMDMs by expressing little to no *Gsdmd* mRNA or GSDMD protein (Figures S1G and S1H). These data suggest that IRF1 and IRF2 engage a single IRF binding site to promote *Gsdmd* expression. Consistent with their lack of GSDMD expression, *Irf1*^-/-^*Irf2^-/-^* and *Gsdmd^Irf/Irf^*BMDMs were as resistant as *Gsdmd^-/-^* BMDMs to LPS-induced pyroptosis (Figures S1I and S1J). *Irf2^-/-^* BMDMs were not as protected as *Irf1*^-/-^*Irf2^-/-^* BMDMs, but died slower than *Irf1^-/-^* BMDMs (Figures S1I and S1J), indicating that IRF2 is the dominant IRF regulating GSDMD expression in BMDMs.

### IRF2 suppresses ISG expression across multiple skin cell types

As reported ^10^, *Irf2^-/-^* mice developed alopecia at around 8 weeks of age, and extensive dermatitis by 12 weeks of age (Figure S2A). Given that *Irf2^-/-^* mice exhibit abnormally high expression of several ISGs ^10^, we performed single cell RNA-seq (scRNA-seq) of *Irf2*^-/-^ skin to investigate which cell type(s) have dysregulated ISG expression. Skin cells were annotated using published marker genes ^20^ and included hair follicle cells, interfollicular epidermal cells, fibroblasts, T cells, macrophages, and Langerhans cells (Figure S2B). WT and *Irf2^-/-^* skin showed similar cell type frequencies, except for macrophages, cycling interfollicular epidermis, and sebaceous gland, which were over-represented in *Irf2^-/-^*skin (Figure S2B).

*Irf1* and *Irf2* were variably expressed across WT skin cells. *Irf2* showed stronger expression in the interfollicular epidermis (Figure S2C), while *Irf1* expression was stronger in basal hair follicle cells and cycling interfollicular epidermal cells (Figure S2C). ISGs were broadly elevated across most *Irf2^-/-^*cell types (Figures S2D and S2E). Interestingly, *Irf2^-/-^* skin macrophages were the most enriched and exhibited the strongest ISG signature of the different cell types (Figures S2D and S2E). Suprabasal interfollicular epidermal cells and hair follicle cells showed the lowest ISG signature, restricted to just a few ISGs like *Isg15* and *Ifi27l2a* (Figures S2D and S2E).

### IRF1 drives elevated ISG expression in the absence of IRF2

Given that IRF1 drove elevated expression of *Casp11* in *Irf2^-/-^* BMDMs (Figures S1A and S1B), we checked whether IRF1 contributed to ISG up-regulation in *Irf2^-/-^*tissues. Overall, IRF1 loss normalized expression of all 296 ISGs elevated in *Irf2*^-/-^ skin (Figure 1A). Importantly, *Irf1*^-/-^*Irf2^-/-^* mice did not develop skin lesions like *Irf2^-/-^* mice (Figures 1B and 1C). Immunolabelling of ISG-encoded ZBP1 revealed that *Irf2^-/-^*skin contained more immune, endothelial, and epithelial cells that were strongly positive for ZBP1 than WT skin, but *Irf1*^-/-^*Irf2^-/-^* skin did not (Figure 1D). These data indicate that IRF1 is the main driver of ISG expression and dermatitis in the absence of IRF2.

**Figure 1.**
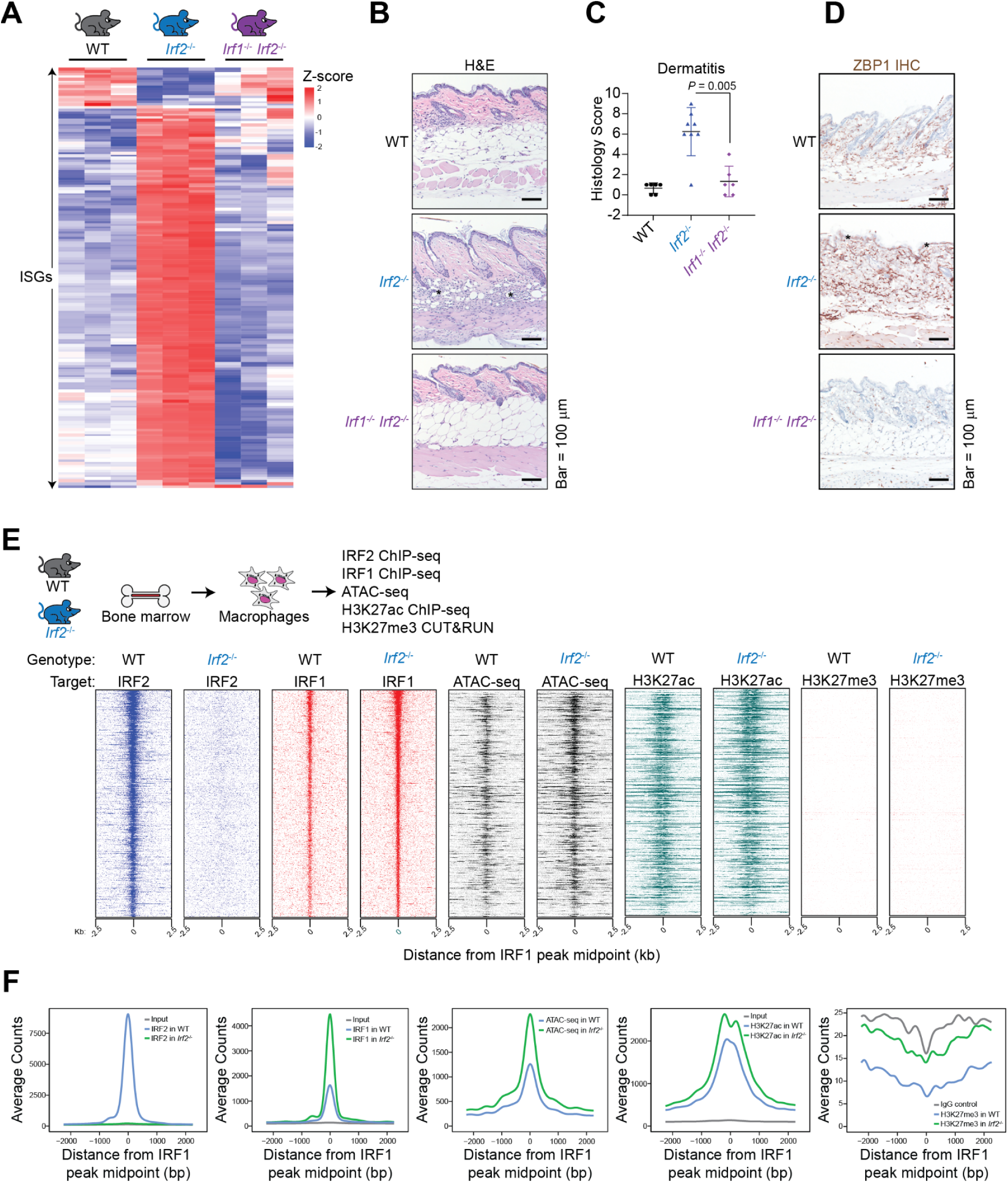
IRF1 drives elevated ISG expression in the absence of IRF2. (A) Heat-map shows gene expression (Z-score) for 327 differentially expressed ISGs (adjusted *P* <0.05) in the skin of mice aged 6 weeks (*n* = 3 mice per genotype). (B) Hematoxylin and eosin-stained skin sections of mice aged 12 weeks. *n* = 5 per genotype. (*) denotes immune infiltrates. Scale bar, 100 μm. (C) Skin histology scores of mice aged 12 weeks. (WT, *n* = 6; *Irf2*^-/-^, *n* = 8; *Irf1*^-/-^*Irf2*^-/-^, *n* = 6). *P* values calculated by two-tailed Mann–Whitney test. (D) ZBP1 immunolabelling (brown) of the skin sections in (B). Asterisk denotes epidermal ZBP1 labelling, which is absent in the other genotypes. (E) IRF2 ChIP-seq, IRF1 ChIP-seq, ATAC-seq, H3K27ac ChIP-seq, and H3K27me3 CUT&RUN in WT and *Irf2*^-/-^ BMDMs. Plots show the distribution of normalized read counts in a 5,000 bp window (binned at 10 bps) around 725 IRF1 peaks in the *Irf2*^-/-^ sample. All rows are linked and sorted by total reads in a 1,000 bp region around IRF1 peaks in the *Irf2*^-/-^ sample. (F) Distribution of gene-averaged read counts in (E). See also Figure S3 and S4.

Skin inflammation in *Irf2^-/-^* mice was accompanied by alterations in the spleen. Marginal zones were prominent in *Irf2^-/-^*spleens and there was increased extramedullary haematopoiesis (Figure S3A). RNA-seq indicated dysregulated ISG expression in *Irf2^-/-^* spleens, although the ISG signature was weaker than in *Irf2^-/-^* skin (Figure S3B). Both the white pulp alterations and elevated ISG expression in *Irf2^-/-^*spleens were normalized by *Irf1* loss (Figures S3A and S3C), indicating that IRF2 limits IRF1 activity in the spleen. Given that *Irf9* deficiency also prevents skin inflammation in *Irf2*^-/-^ mice ^10^, we speculate that IRF1 and IRF9 both contribute to ISG expression and have non-redundant roles acting at IRF regulatory elements and ISREs, respectively ^21^.

### Chromatin profiling reveals that IRF2 functions as a suppressor exclusively at IRF1 binding sites

Next we investigated how IRF2 activates genes like *Gsdmd* ^18^ and *Tlr3* ^16,17^, but represses genes like *Casp11* (Figures S1A). Macrophages showed the strongest ISG signature in skin (Figures S2D and S2E), so we focussed on this cell type for further mechanistic studies. ChIP-seq of WT and *Irf2^-/-^*BMDMs revealed 725 IRF1 binding sites, predominantly near innate immune genes (Figures 1E, 1F, and S4A). In WT cells, IRF2 was detected at 676 (93%) of these IRF1 binding sites (Figures 1E and 1F). Of these 676 sites, 496 mapped to enhancers (> 2 kb from transcription start sites [TSS]), while 180 peaks were at ISG promoters, including those of *Gbp2*, *Ifit1*, and *Zbp1* (Figure S4B). IRF2 loss at these sites coincided with increased H3K27ac, the mark of active promoters and enhancers, and the chromatin was more open than in WT BMDMs by assay for transposase-accessible chromatin (ATAC)-seq (Figures 1E, 1F, and S4B). Consistent with IRF1 driving ISG expression in *Irf2^-/-^* cells (Figure 1A), more chromatin opening in *Irf2*^-/-^ BMDMs coincided with increased IRF1 binding at these sites (Figures 1E and 1F).

We used native gel shift assays to examine co-binding of IRF1 and IRF2 to DNA. A DNA probe consisting of a *Gpb2* promoter sequence with a single AAA_NN_GAAA_NNN_AAA motif was shifted by purified IRF1 or IRF2 (Figure S4C). Mutating the GAAA core and the 5’-flanking AA sequence ^13,14^ abrogated IRF1 and IRF2 binding, confirming the specificity of the assay. An IRF1-IRF2-DNA complex was detected when adding IRF1 to IRF2-DNA complexes, and required the 3’-flanking AAA sequence in the IRF motif (Figure S4D). Since IRF1 was unable to bind to DNA with a single AAA core (Figure S4C), our data suggests a degree of cooperativity, rather than competition, between IRF1 and IRF2 in DNA binding.

Despite IRF2 suppression of ISG expression, IRF2 binding was not associated with H3K27me3 or H3K9me3 histone modifications, markers of facultative and constitutive repressive heterochromatin, respectively (Figures 1E and 1F). IRF2 binding genome-wide was only associated with the active transcription marker H3K27ac (Figure S4E). Furthermore, *Irf2*-deficiency did not alter the distribution of repressive H3K27me3 across TSS genome-wide (Figure S4F). IRF2 interacts with BRD7, a component of the SWI/SNF chromatin remodeling complex ^22,23^, and this interaction might facilitate opening of the chromatin for IRF2-mediated *Tlr3* transcription ^16^. This possibility prompted us to compare WT and *Irf2*^-/-^ BMDMs in their utilization of SMARCA4, a core component of the SWI/SNF complex ^24^. IRF2 loss altered the distribution of SMARCA4 at 1216 sites genome-wide (Figures S4G) and reduced SMARCA4 occupancy at 1019 (84%) of these sites. These data suggest that IRF2 largely acts as a transcriptional activator. Sites with increased SMARCA4 occupancy were enriched with IRF1 binding sites (Figure S4G). These data indicates that IRF1 drives the increase of SMARCA4 occupancy in the absence of IRF2.

Examining the expression of genes ‘activated’ or ‘suppressed’ by IRF2, we found that WT skin expressed the ‘suppressed’ genes, on average, as much as the ‘activated’ genes (Figure S4H). In other words, IRF2 did not silence the ‘suppressed’ genes. Rather, these genes were transcriptionally active and their expression was amplified by IRF1 once IRF2 was lost (Figures S4I).

### Innate immune signaling triggers IRF2 degradation

Next, we assessed the role of IRF1 in ISG induction by different stimuli. Consistent with prior studies ^8, 9^, *Irf1^-/-^* BMDMs exhibited less ISG transcription than WT BMDMs after treatment with IFN-γ (Figure S5A) or TLR4 agonist LPS (Figure 2A), whereas IRF1 was largely dispensable for ISG induction by IFN-□ (Figure 2B). Only 17 of the 127 ISGs induced in WT cells by IFN-□ after 4 hours showed no induction in *Irf1*^-/-^ BMDMs. More IFN-□-induced ISG transcripts were abrogated in *Irf9^-/-^* BMDMs, but it was the combined loss of IRF1 and IRF9 that suppressed ISG transcription the most (Figure 2B). These data suggest that IRF9 is the primary activator of IFN-β-induced ISGs, but IRF1 contributes to ISG activation when IRF9 is lost. IRF1 contributed to ISG transcription downstream of multiple TLRs, not just TLR4, because *Irf1^-/-^* BMDMs also exhibited defective ISG transcription in response to TLR3 agonist poly(I:C) and TLR1/2 agonist Pam3CSK4 (Figure S5A).

**Figure 2.**
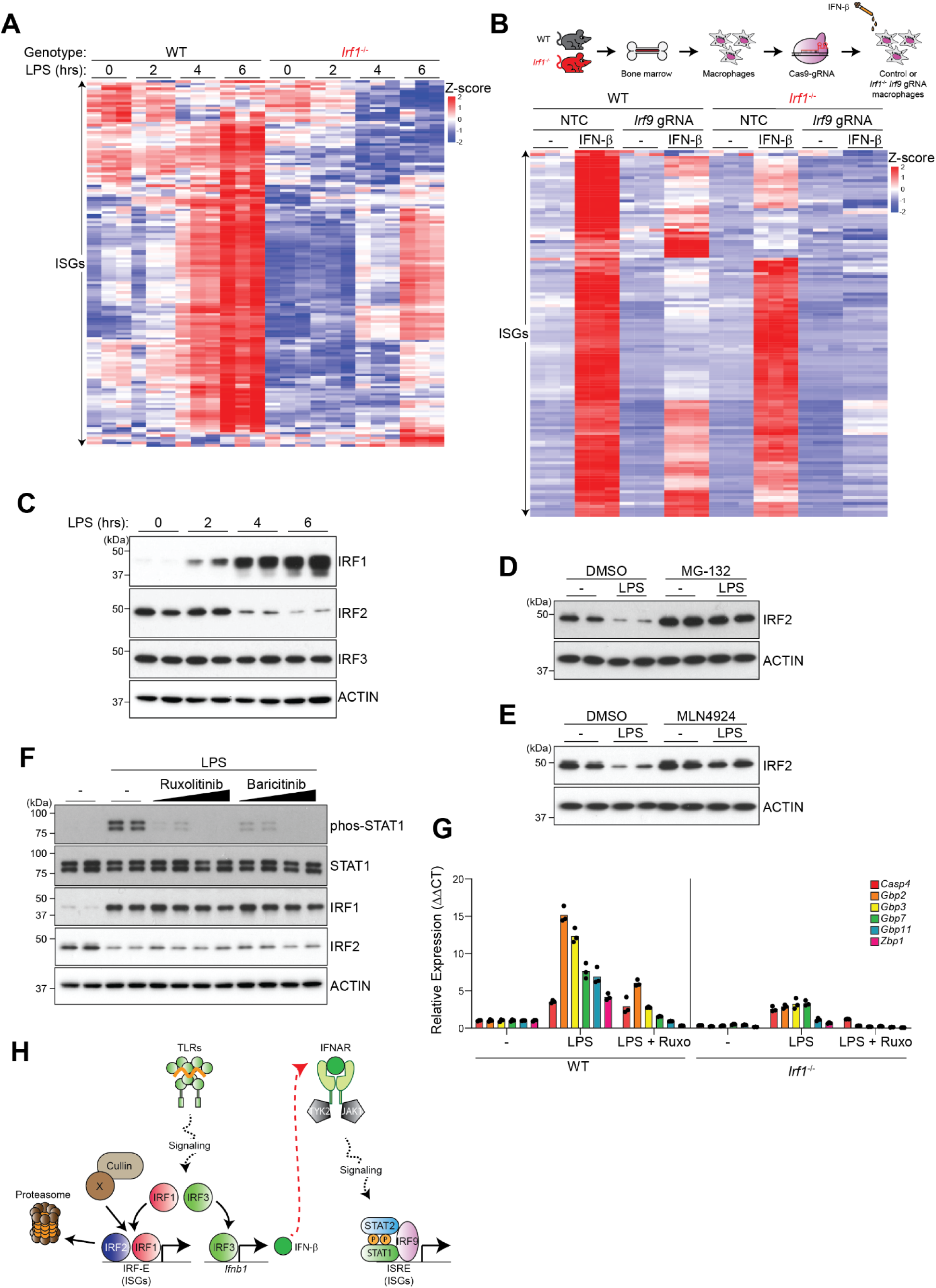
TLR signaling triggers proteasomal degradation of IRF2. (A) Heat-map shows gene expression (Z-score) for 189 differentially expressed ISGs (adjusted *P* <0.05) in LPS-treated BMDMs (*n* = 3 mice per genotype). (B) Heat-map shows gene expression (Z-score) for 127 differentially expressed ISGs (adjusted *P* <0.05) in BMDMs treated with IFN-□ for 4 hours (*n* = 3 mice per genotype). (C) Immunoblots of BMDMs (*n* = 2 per genotype). (D and E) Immunoblots of BMDMs treated with MG-132 (D) or MLN4924 (E) for 15 minutes, and then stimulated with LPS for 4 hours (*n* = 2 per genotype). (F) Immunoblots of BMDMs treated with Ruxolitinib and Baricitinib for 15 minutes, and then stimulated with LPS for 4 hours (*n* = 2 per genotype). (G) RT-qPCR of ISGs in BMDMs treated with Ruxolitinib for 15 minutes and then LPS for 4 hours. Mean expression levels of each gene are shown relative to WT BMDMs without treatment. Circles, BMDMs from individual mice. Results representative of 3 independent experiments. (H) Model for ISG transcription in LPS-treated cells. IRF1 contributes to ISG expression independent of IFN signaling. See also Figure S5.

To investigate how IRF1 overcomes IRF2 interference following TLR engagement, we monitored IRF levels after LPS stimulation. WT BMDMs expressed more *Irf1* mRNA and IRF1 protein after LPS treatment, whereas IRF2 decreased and IRF3 was unchanged (Figure 2C and S5B). *Irf2* mRNA abundance was unchanged, suggesting LPS might promote IRF2 protein degradation (Figure S5B). The proteasome inhibitor MG-132 (Figure 2D) or MNL4924, a NEDD inhibitor that blocks the activity of Cullin E3 ubiquitin ligases (Figure 2E), prevented LPS-induced IRF2 loss in BMDMs, indicating that IRF2 is targeted by a Cullin E3 ubiquitin ligase for proteasomal degradation. IFN-γ, poly(I:C), Pam3CSK4, and the TLR7 agonist Imiquimod also caused IRF2 loss in BMDMs (Figure S5C), suggesting that this is a universal mechanism for unleashing IRF1-driven ISG transcription.

To determine if LPS-induced IRF2 loss requires paracrine/autocrine signaling triggered by IFN secretion, we used Ruxolitinib or Baricitinib to inhibit JAK-STAT signaling. Both compounds prevented STAT1 phosphorylation in LPS-stimulated BMDMs without affecting the induction of IRF1 or degradation of IRF2 (Figure 2F). Interestingly, Ruxolitinib reduced LPS-induced ISG expression in WT and *Irf1^-/-^* BMDMs (Figures 2G), suggesting that IRF1 contributes to ISG expression independent of the paracrine/autocrine IFN pathway. Deletion of *Irf9* or *Ifnar1* also had no impact on IRF1 expression or IRF2 degradation in response to LPS, but when combined with *Irf1* deficiency, further suppressed LPS-induced ISG transcription (Figures S5D-S5G). Transcription factor PRDM1 (also called BLIMP-1) may repress IRF1 and IRF2 activity by competing for the same DNA binding motif ^25^. We confirmed PRDM1 induction in LPS-treated BMDMs, but *Prdm1* deletion did not affect the induction of IRF1, degradation of IRF2, or ISG transcription (Figures S5H and S5I). Collectively, our data suggest that ISG transcription in LPS-treated BMDMs is driven by two distinct pathways: an early pathway initiated by IRF1, and a secondary pathway triggered when secreted IFN-α/□ activates IFNAR signaling (Figure 2H).

### IRF9 transcribes *Irf2* during IFN-□ stimulation, countering its degradation

In contrast to the TLR agonists and IFN-γ, IFN-□ induced IRF1 in BMDMs without overt IRF2 loss (Figure S6A). Despite this different pattern of IRF regulation, LPS and IFN-□ induced comparable ISG expression (Figure S6B). IFN-□ differed from LPS in stimulating IRF9-dependent, IRF1-independent transcription of *Irf2* (Figure S6C). IRF9 was bound to the *Irf2* promoter after IFN-□ stimulation and this coincided with increased H3K27ac, the marker of active transcription (Figure S6D). When *Irf9* was deleted from BMDMs to prevent *Irf2* transcription, IFN-□ stimulated IRF2 loss that was prevented by MNL4924 (Figures 3A and 3B). Therefore, IRF2 is degraded in the proteasome after IFN-□ stimulation, but IRF9 increases *Irf2* transcription to maintain IRF2 at steady state levels. Note that IFN-□ did not cause IRF2 loss in BMDMs lacking both IRF1 and IRF9 (Figure 3A). The role of IRF1 in IRF2 degradation is investigated further below.

**Figure 3.**
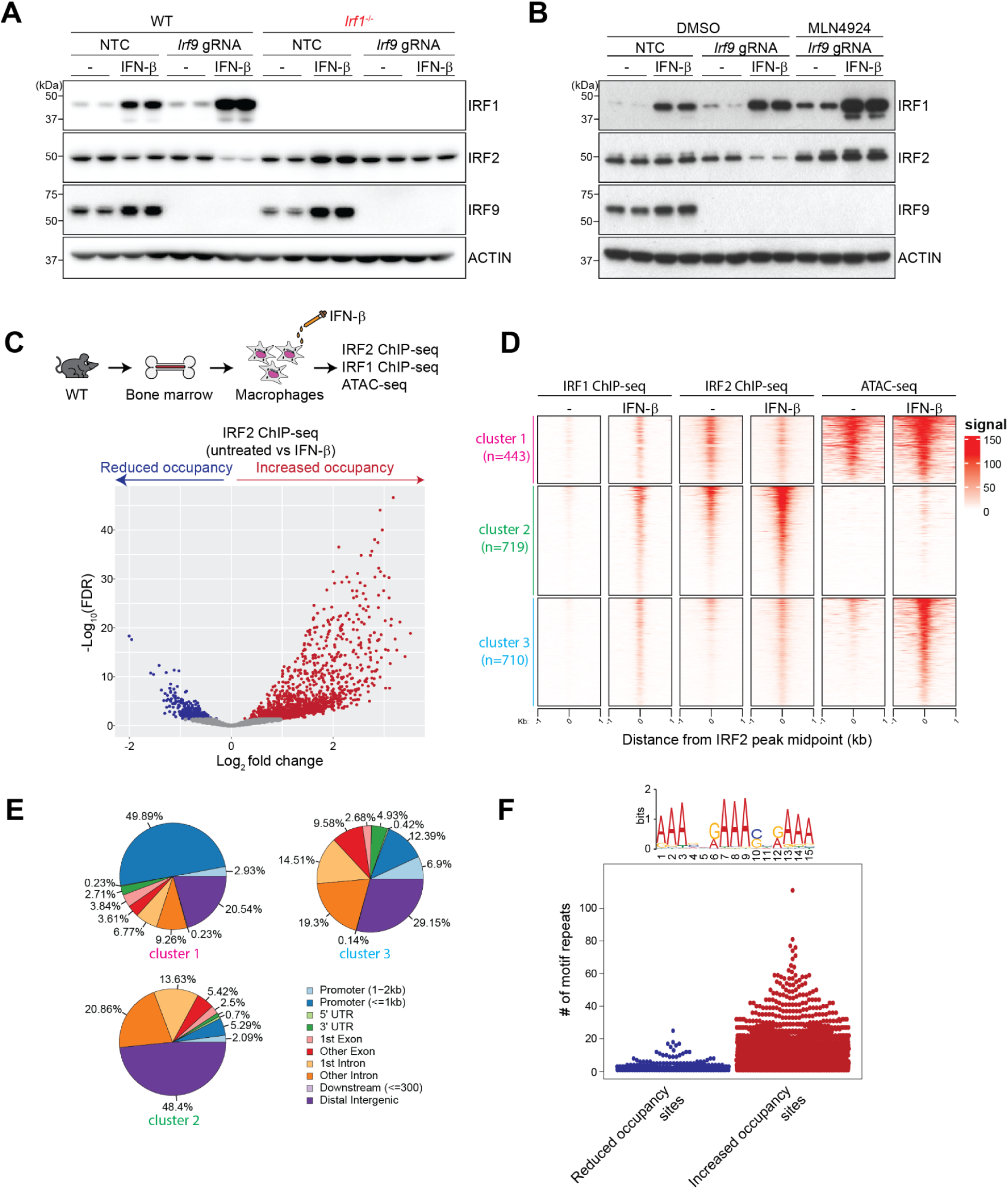
IRF9-driven *Irf2* transcription masks IRF2 protein loss during IFN-□ stimulation. (A) Immunoblots of BMDMs treated with IFN-□ for 4 hours (*n* = 2 per genotype). (B) Immunoblots of BMDMs treated with MLN4924 for 15 minutes and then IFN-□ for 4 hours (*n* = 2 per genotype). (C) Volcano plots of IRF2 binding, as measured by ChIP-seq, in BMDMs treated with IFN-□ for 4 hours (*n* = 2 biological replicates using cells from individual mice). Dots represent different IRF2 binding sites. Sites are colored if they meet the following criteria: log2 fold change <-1 or >1 and adjusted *P* <0.05 (Wald test). Grey dots, *P* > 0.05. (D) K-means clustering of IRF1 ChIP-seq, IRF2 ChIP-seq, and ATAC-seq at differentially bound IRF2 sites in (C). Heatmap shows signal intensity across each cluster, centered on IRF2 peaks in the WT untreated sample, with a +/-1 Kb region shown. Results representative of 2 biological replicates using cells from different mice. (E) Piecharts showing distribution of peaks in clusters defined in (D). (F) Top: *de novo* motif search at sites of reduced and increased IRF2 occupancy in (C). Bottom: enrichment of IRF motif repeats at sites of reduced and increased IRF2 occupancy in (C). Dots represent different IRF2 binding sites. See also Figure S6.

Given that IRF1, IRF2, and IRF9 bind to similar DNA motifs within ISG promoters, it is unclear how IRF9 initiates transcription when IRF2 is present. ChIP-seq revealed that IFN-□ stimulation reduced IRF2 occupancy at 443 sites and increased occupancy at 1429 sites in the genome (Figure 3C). K-means clustering of IRF2 ChIP-seq, IRF1 ChIP-seq, and ATAC-seq indicated that these sites fell into 3 groups (Figure 3D). Cluster 1 sites had less IRF2 and more IRF1 bound after IFN-□ stimulation, retained high chromatin opening, and were predominately located within the promoters of innate immune genes, including ISGs (Figures 3D, 3E and S6E). Sites in clusters 2 and 3 exhibited increased IRF1 and IRF2 occupancy after IFN-□ stimulation, but were distinguished by their chromatin opening (Figure 3D). Cluster 2 sites remained largely closed, were not at promoters (5.29% for cluster 2 vs 49.89% for cluster 1), and were not enriched near innate immune genes (Figures 3D, 3E, and S6E). Cluster 3 sites were more open after IFN-□ stimulation, not at promoters (12.39% for cluster 3 vs 49.89% for cluster 1), and enriched near innate immune genes, including ISGs (Figures 3D, 3E, and S6E). Curiously, LPS stimulation increased the accessibility of cluster 2 and 3 sites in an IRF1-dependent manner (Figures S6F and S6G). These data indicate that IRF2, while not eliminated by IFNAR signaling, is redistributed within the genome.

Motif analysis showed that sites in all 3 clusters contained the canonical AAA_NN_GAAA_NNN_AAA IRF motif, but sites in clusters 2 and 3 were in microsatellite loci characterized by abundant NNNAAA repeats. Reduced occupancy cluster 1 sites typically contained 2 to 3 repeats, whereas increased occupancy sites averaged 11 repeats, with some having between 60 to 80 repeats (Figure 3F). Thus, IFN-□ stimulates chromatin opening at sites distal from ISG promoters and containing multiple IRF motifs engaged by IRF2.

### IRF1 triggers SPOP-mediated degradation of IRF2

To identify the Cullin E3 ubiquitin ligase(s) responsible for IRF2 degradation, we determined if a specific member of the Cullin family was required. *Cul3* deletion in BMDMs prevented IRF2 loss after LPS stimulation, whereas deletion of *Cul1*, *Cul2*, *Cul4a*, *Cul4b* or *Cul5* did not (Figure S7A). Cullin-3 binds to more than 100 different substrate adaptors ^26^, but not all of them are expressed in macrophages. RNA-seq identified several Cul3 substrate adaptors expressed in BMDMs, including *Lgals3bp*, *Rcbtb2*, and *Spop* (Figure S7B). *Spopl* shares 82% sequence identity with *Spop* ^27^, but was not expressed in BMDMs (Figure S7B). We deleted 18 of the most expressed Cul3 substrate adaptors in BMDMs and only *Spop* deletion prevented IRF2 loss after LPS (Figures 4A and S7C) or IFN-□ stimulation (Figures S7D).

**Figure 4.**
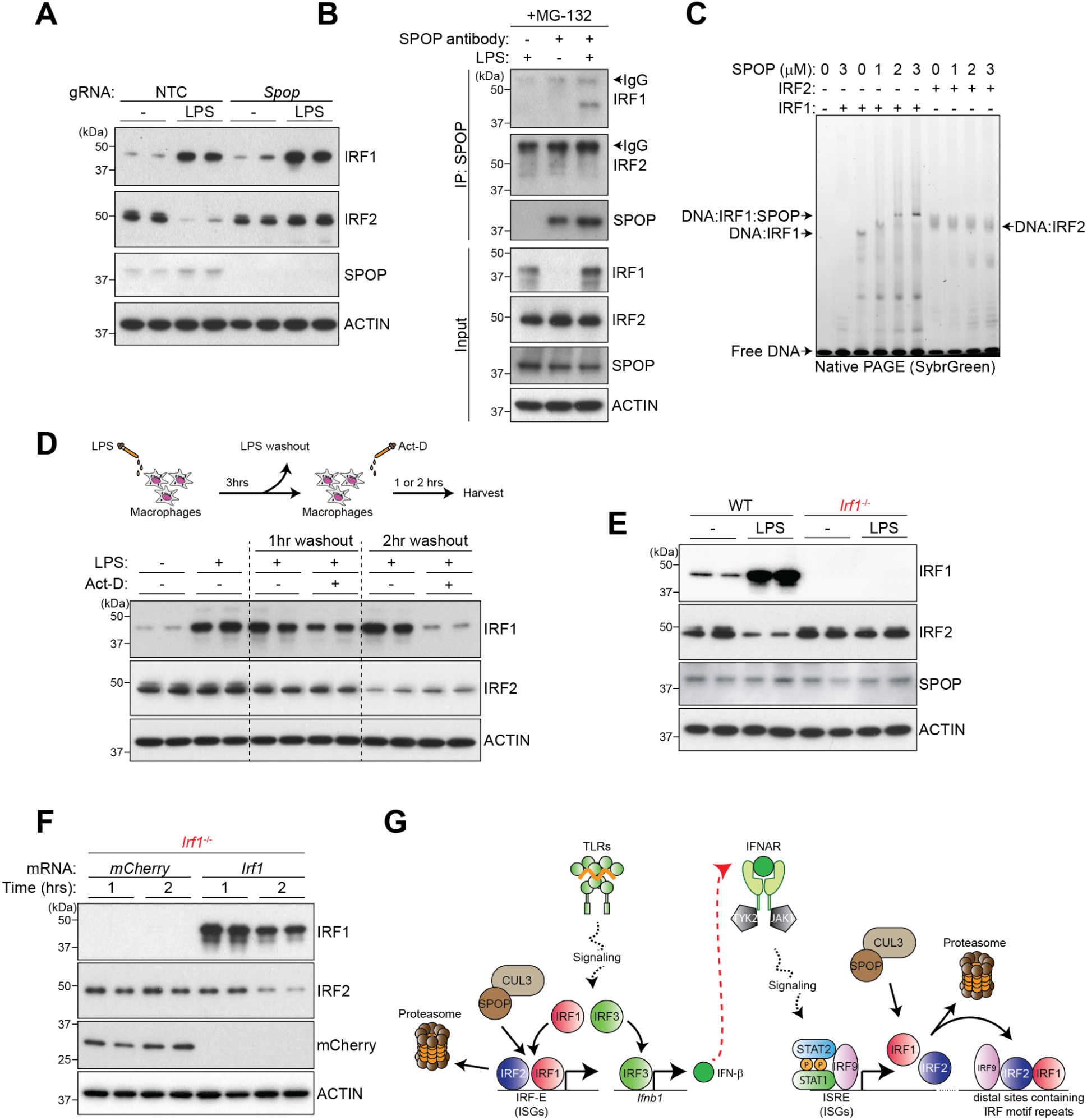
IRF1 is required for SPOP to target IRF2. (A and E) Immunoblots of BMDMs treated with LPS for 4 hours. Lanes represent BMDMs from individual mice (*n* = 2 per genotype). (B) Immunoblots after SPOP immunoprecipitation (IP) from BMDMs treated with MG-132 and LPS for 4 hours. Results representative of 2 independent experiments. (C) Gel shift assay of recombinant IRF1 and IRF2 (0.5 uM each) bound to a DNA sequence from the *Gbp2* promoter in the presence of recombinant SPOP. Gels were stained with SybrGreen to visualize complexes. (D) Immunoblots of BMDMs treated with LPS for 3 hours, and then actinomycin-D for 2 hours. Lanes represent BMDMs from individual mice (*n* = 2 per genotype). (F) Immunoblots of BMDMs electroporated with *Irf1* or *mCherry* mRNA. Lanes represent BMDMs from individual mice (*n* = 2). Results representative of 3 independent experiments. (G) Model of ISG induction in LPS-stimulated BMDMs. IRF1 induction drives early ISG transcription and facilitates the degradation of IRF2 by SPOP. See also Figure S7.

IRF1 co-immunoprecipitated with SPOP in LPS-stimulated BMDMs, but IRF2 did not (Figure 4B). This result is consistent with SPOP targeting IRF1 in cancer cell lines ^28,29^. In native shift assays, recombinant SPOP did not bind to the DNA probe on its own, but appeared to shift some of the IRF1-DNA complex, but not the IRF2-DNA complex, at higher concentrations (Figure 4C). SPOP may bind to IRF1 rather than IRF2 because IRF1 contains one canonical and two substandard SPOP binding motifs, while IRF2 has a single substandard SPOP binding motif (Figure S7E). The SPOP-IRF1-DNA complex also required high concentrations of IRF1 (Figures S7F). The interaction between IRF1 and SPOP prompted us to test if IRF1 is degraded after LPS stimulation, but replaced by newly synthesized IRF1 as *Irf1* transcription increases. The transcriptional inhibitor actinomycin D (Act–D) caused IRF1 loss in BMDMs pre-stimulated with LPS, confirming that IRF1, like IRF2, is labile after LPS stimulation (Figure 4D).

Reminiscent of our earlier observation in BMDMs lacking *Irf1* and *Irf9*, which failed to degrade IRF2 in response to IFN-□ (Figure 3A), *Irf1*^-/-^ BMDMs did not lose IRF2 after LPS stimulation despite expressing SPOP (Fig. 4E). We propose that IRF1 brings SPOP to genomic sites occupied by IRF2 thereby promoting IRF2 degradation. Consistent with this model, *Irf1*^-/-^ BMDMs reconstituted with *Irf1* mRNA, but not *mCherry* mRNA, exhibited robust IRF1 expression after 1 hour, followed by a drop in IRF2 at 2 hours (Figure 4F). Thus, IRF1 is sufficient for IRF2 degradation and other signaling events are not required. SPOP was required for IRF2 degradation because *Irf1* mRNA delivery into *Spop*-deficient BMDMs did not cause IRF2 loss, despite increased IRF1 expression (Figure S7G). Collectively, our data indicate that increased IRF1 expression triggered by TLR or IFNAR signaling promotes SPOP-dependent IRF2 degradation (Figure 4G).

### SPOP prevents aberrant expression of proinflammatory genes by targeting IRF2 for degradation

Sustained IRF2 expression in *Spop*-deficient BMDMs might limit IRF1-driven ISG expression or cause aberrant gene expression. Consistent with the TLR adaptor protein MYD88 being an SPOP substrate ^30,31,32,33^, *Spop-*deficient BMDMs expressed more MYD88 than control BMDMs, but this did not impact LPS-induced IKK and TBK1 phosphorylation or Iκ□α degradation (Figure S8A and S8B). ChIP-seq indicated that LPS stimulation of *Spop*-deficient BMDMs altered the distribution of IRF2 at 242 sites genome-wide, 224 (92%) of them showing increased IRF2 occupancy (Figures 5A and S8C). Containing the canonical IRF motif, these sites were located in distal intergenic regions, predominantly near genes involved in chemokine signaling rather than ISGs (Figures S8D, 5B and 5C). RNA-seq of control and *Spop*-deficient BMDMs after LPS stimulation indicated differential expression of 124 genes (*P* = 0.05, Log2 fold-change < or > 1.5; Figure 5D). To determine if the elevated gene expression in *Spop*-deficient BMDMs was due to IRF2, we deleted *Spop* in WT and *Irf2*^-/-^ BMDMs and stimulated the cells with LPS for 0, 6, or 24 hours.

**Figure 5.**
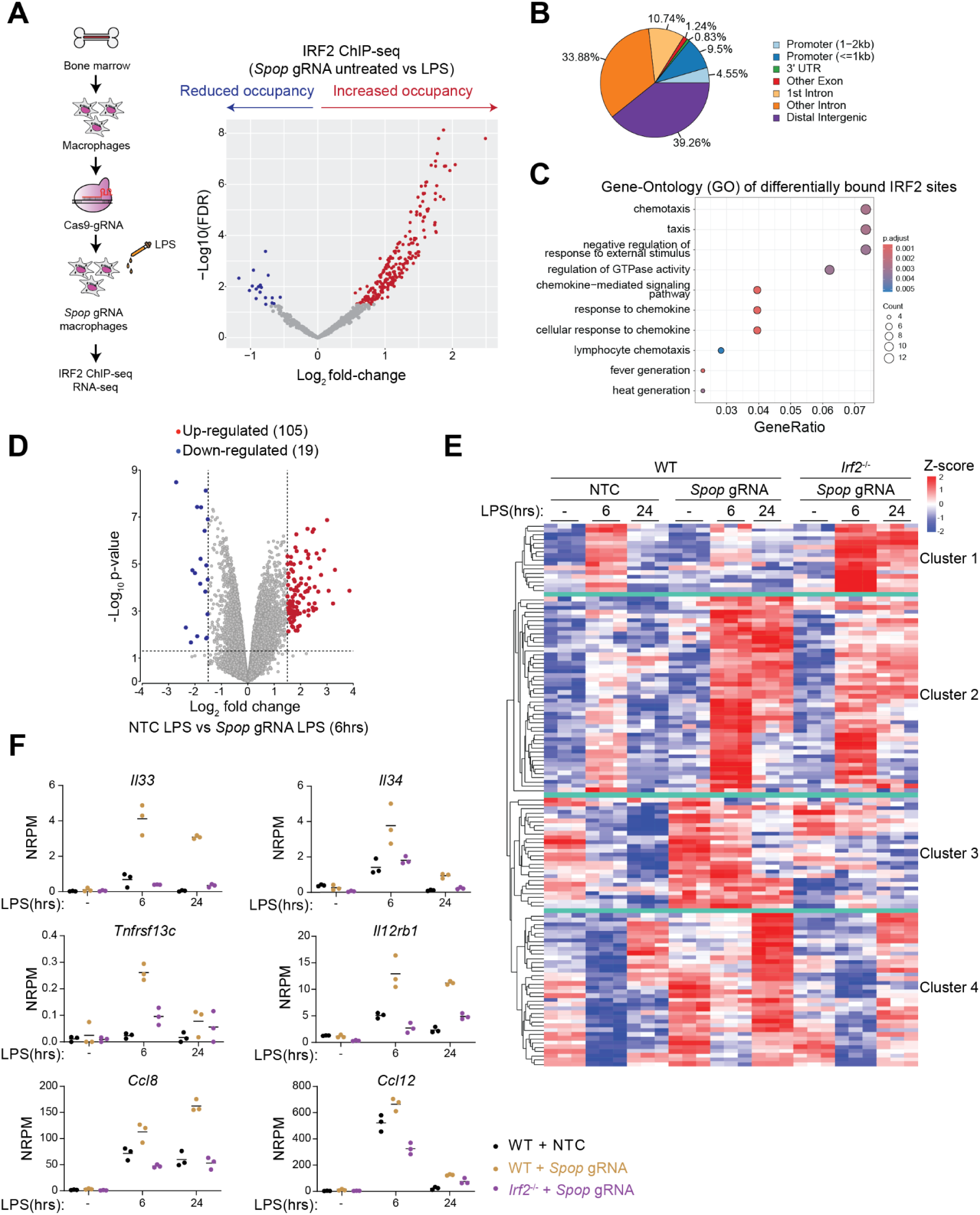
IRF2 drives aberrant gene transcription in the absence of SPOP. (A) Volcano plot shows changes in IRF2 binding, as measured by ChIP-seq, in BMDMs treated with LPS for 4 hours (*n* = 2 biological replicates using cells from individual mice). Dots represent different IRF2 binding sites. Sites are colored if they meet the following criteria: log2 fold change <-1 or >1 and adjusted *P* <0.05 (Wald test). Grey dots, *P* > 0.05. (B) The distribution of sites with significantly increased (log2 fold change <-1 and adjusted *P* <0.05) IRF2 occupancy in (A). (C) Enriched ontology terms for significantly increased (log2 fold change <-1 and adjusted *P* <0.05) IRF2 occupancy peaks in (A). (D) Volcano plot shows changes in gene expression in BMDMs treated with LPS for 4 hours (*n* = 3 biological replicates using cells from individual mice). Genes are colored if they meet the following criteria: log2 fold change <-1.5 or >1.5 and adjusted *P* <0.05. Grey dots, *P* > 0.05. (E) Heat-map shows gene expression (Z-score) for 124 differentially expressed genes (adjusted *P* <0.05, log2 fold change <-1.5 or >1.5) in BMDMs treated with LPS (*n* = 3 mice per genotype). (F) Expression of selected cytokine– and chemokine– related genes in LPS-treated BMDMs. NRPKM, normalized reads per kilobase per million reads. Symbols, BMDMs from individual mice (*n* = 3 per genotype). Lines indicate the mean. See also Figure S8.

Genes differentially expressed in *Spop*-deficient BMDMs fell into 4 clusters (Figure 5E). Cluster 1 genes exhibited impaired induction in the absence of *Spop,* were largely normalized in cells lacking *Irf2* and *Spop*, and included several ISGs that are IRF1 target genes (Figures 5E, S8E, and S8F). Cluster 2 genes were transiently induced in control BMDMs, but exhibited a sustained elevation in *Spop*-deficient BMDMs (Figure 5E). Their expression decreased in BMDMs lacking *Spop* and *Irf2*, indicating that IRF2 partly contributes to their transcription (Figure 5E). Genes in clusters 3 and 4 were elevated in *Spop*-deficient BMDMs, even without LPS stimulation, and were distinguished by differences in their kinetics of induction. Cluster 3 and 4 genes were largely normalized in *Spop*–deleted *Irf2*^-/-^ macrophages (Figures 5E and 5F), indicating that IRF2 regulates their expression. Gene-Ontology analysis of the genes in clusters 2, 3, and 4 indicated enrichment in serine peptidase activity, plus cytokine and chemokine signaling (Figure S8G), Together, our ChIP-seq and RNA-seq datasets indicate that SPOP-driven IRF2 degradation prevents aberrant expression of proinflammatory genes.

### SPOP targets IRF2 at cytokine loci

*Spop* deficient mice die shortly after birth ^34^, so we validated our results obtained with CRISPR/Cas9 gene editing by using *Spop^-/-^* BMDMs from tamoxifen-treated *Spop^fl/fl^ Rosa26*^CreERT^^2/+^ mice. *Spop^-/-^* BMDMs lacked detectable SPOP and failed to degrade IRF2 when challenged with formaldehyde-inactivated *E. Coli* or *S. Aureus* (Figure 6A). *Spop*^-/-^ BMDMs also expressed more IRF1, consistent with it being an SPOP substrate (Figure 6A). Stimulated *Spop*^-/-^ BMDMs released more CCL2, CCL7, CXCL1, and IL-22 than their WT counterparts, but released normal amounts of IL-6 and TNF (Figures 6B and S9A). Thus, SPOP loss only impacts select cytokines and chemokines.

**Figure 6.**
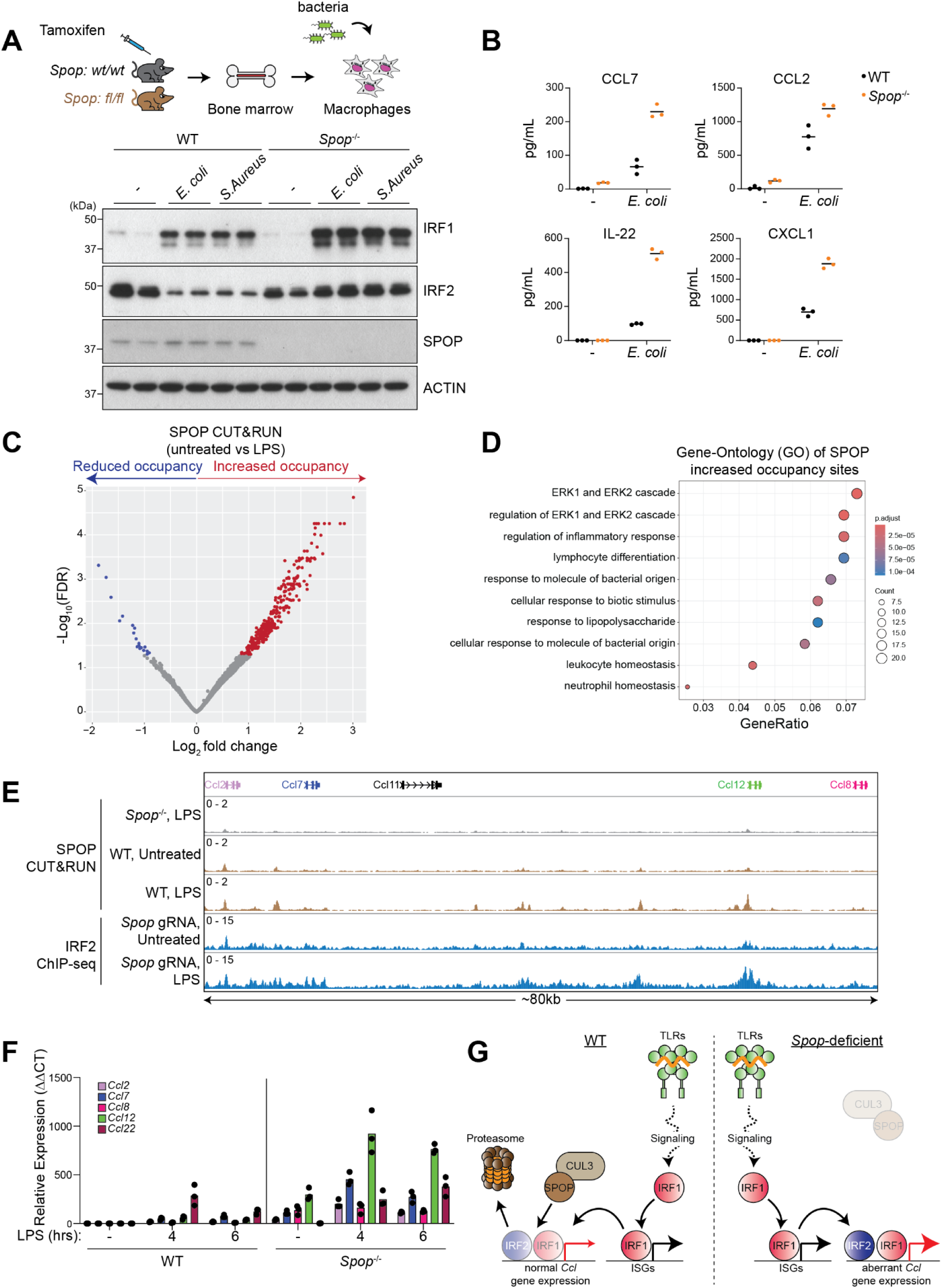
IRF1 is required for IRF2 degradation. (A) Immunoblots of BMDMs treated with formaldehyde-inactivated *E. coli* or *S. Aureus* particles for 4 hours (*n* = 2 per genotype). (B) CCL7, CCL2, IL-22, and CXCL1 secreted from BMDMs in (A). Circles, cells from individual mice. Lines, the mean. (C) Volcano plot shows changes in SPOP binding, as measured by CUT&RUN, in BMDMs treated with LPS for 4 hours (*n* = 2 biological replicates using cells from individual mice). Dots represent different SPOP binding sites. Sites are colored if they meet the following criteria: log2 fold change <-1 or >1 and adjusted *P* <0.05 (Wald test). Grey dots, *P* > 0.05. (D) Enriched ontology terms for significantly increased (log2 fold change <-1 and adjusted *P* <0.05) SPOP peaks in (C). (E) SPOP CUT&RUN and IRF2 ChIP-seq at the *Ccl* loci in BMDMs treated with LPS for 4 hours. Numbers denote track scales (normalized read counts). Results representative of 2 biological replicates using cells from individual mice. (F) RT-qPCR of cytokines in BMDMs treated with LPS. Mean expression levels of each gene are shown relative to those in untreated WT BMDMs. Dots, BMDMs from individual mice. Results representative of 3 independent experiments. (G) Model for IRF2 driving cytokine and chemokine gene expression in *Spop*-deficient BMDMs. See also Figure S9.

Cleavage Under Targets and Release Using Nuclease (CUT&RUN) analysis revealed 4422 SPOP genomic binding sites (Figure S9B). Importantly, enrichment for these sites decreased in *Spop*^-/-^ BMDMs. The sites were predominantly near genes involved in immune receptor signaling and myeloid cell differentiation (Figure S9C). LPS stimulation increased SPOP occupancy at 316 genomic sites (Figure 6C) and gene-ontology analysis revealed terms related to the LPS response and ERK signaling (Figure 6D). To identify genomic loci where SPOP potentially targets IRF2, we compared our SPOP CUT&RUN data with our earlier IRF2 ChIP-seq data in *Spop*-deficient BMDMs. K-means clustering identified overlapping SPOP and IRF2 signals at 207 (66%) of the sites exhibiting increased SPOP occupancy (Figures S9D and S9E). These sites were enriched near cytokine- and chemokine-related loci, including *Ccl12* and *Cxcl10*, but also at genes involved in proinflammatory signaling like *Nfkbia* and *AW112010* ^35^ (Figure S9E). We noted increased SPOP binding near several *Ccl* genes in WT BMDMs exposed to LPS (Figure 6E). SPOP loss coincided with increased IRF2 binding and elevated transcription of *Ccl2*, *Ccl7*, *Ccl8*, *Ccl12*, and *Ccl22* (Figures 6E and 6F). LPS-treated *Irf2*^-/-^ BMDMs had decreased expression of these cytokines, consistent with IRF2 regulating their transcription (Figure S9F). Interestingly, LPS-stimulated *Irf1*^-/-^ BMDMs also exhibited less *Ccl2* and *Ccl7* expression than control BMDMs, suggesting that IRF1 and IRF2 cooperate in regulating their expression (Figure S9F). By contrast, *Ccl5* expression only required IRF1 (Figure S9F). Our data suggest that SPOP regulates transcription factors, including IRF1 and IRF2, to prevent aberrant transcription of select cytokines and chemokines (Figure 6G).

## Discussion

ISGs are crucial for controlling bacterial and viral infections. However, ISG expression must be tightly regulated because uncontrolled ISG expression can cause tissue damage. ISG expression triggered by the canonical JAK/STAT/ISGF3 pathway downstream of IFN-α/□ and IFN-γ is well studied, so we were surprised to find that IRF1 drives ISG expression independent of JAK-STAT signaling. IRF1 and IRF2 were the first IRF family members described, but have been studied less than IRF3 and IRF9. IRF2 regulates several genes critical for innate immune defense, including *Gsdmd* and *Tlr3*, but it also limits IRF1-mediated ISG expression. Our data indicates that IRF2 is not a transcriptional repressor. Instead, its apparent suppression of ISGs stems from it being a weaker transcriptional activity than IRF1, which binds to many of the same genomic sites as IRF2. Consequently, IRF2 loss favors IRF1-driven ISG transcription.

Given that IRF1 mediates innate immune defense against mycobacterial infections ^8^ and was needed for robust ISG transcription downstream of several TLRs, we investigated how cells switch from using IRF2 to using IRF1. Expression of IRF1 and IRF2 was regulated by TLR signaling independent of the canonical IFN pathway, with IRF2 being degraded as IRF1 was induced. We propose that IRF2 degradation promotes rapid IRF1-driven ISG expression, and the secretion of IFNs amplifies ISG expression by activating IRF9. However, IRF1 does bind to ISG promoters and enhancers in cells stimulated with IFN-□. We speculate that the relative importance of IRF1 and IRF9 in different settings reflects the temporal sequence of IRF activation. Cells exposed to TLR ligands transcribe both *Irf1* and *Ifnb1*. Consequently, IRF1 can initiate ISG transcription prior to, or in conjunction with, IFN-□ activating IFNAR and IRF9.

A notable finding of our study is that IRF2 degradation required IRF1 and the E3 ubiquitin ligase SPOP. Although we failed to detect SPOP interacting with IRF2, IRF1 recruited SPOP to sites occupied by IRF2, so it is conceivable that transient or indirect interactions then allowed SPOP to ubiquitylate IRF2. Interestingly, cells exposed to IFN-□ degraded IRF2, but this was masked by IRF9-driven *Irf2* transcription. In this setting, IRF2 moved from ISG promoters to distal enhancers containing expanded IRF motif repeats. In effect, relocalization of IRF2 was dominant over IRF2 degradation in promoting transcription by IRF1 and IRF9. Different outputs of a transcription factor at distinct genomic sites have also been reported for EWS-FLI1 in Erwing sarcoma ^36,37,38,39^. EWS-FLI1 multimerizes at GGAA repeats to promote chromatin opening and generate de novo enhancers, but it also inactivates enhancers containing canonical ETS motifs by displacing ETS transcription factors ^36,37,38,39^.

Unexpectedly, *Spop*-deficient macrophages secreted more cytokines and chemokines after LPS stimulation than control cells. The absence of SPOP at promoters and enhancers of these genes coincided with increased IRF2 at these sites. *Irf1*^-/-^ BMDMs, which also failed to degrade IRF2, failed to transcribe the same chemokines. Considering that SPOP targets IRF1 and IRF2, we speculate that the disparity between *Spop*- and *Irf1*-deficient BMDMs arises from the retention of both IRF1 and IRF2 in the former, and only IRF2 in the latter. IRF2 contributed to dysregulated gene expression in *Spop*-deficient BMDMs, but we note that several cluster 2 genes remained elevated in BMDMs lacking both *Spop* and *Irf2*. We speculate that these genes are regulated by IRF1. Alternatively, they may be AP–1 targets because SPOP CUT&RUN peaks were enriched for ERK-related sites. Overall, our findings suggest that SPOP targeting of IRF1 and IRF2 during LPS sensing mitigates aberrant cytokine expression.

---

**Figure S1.**
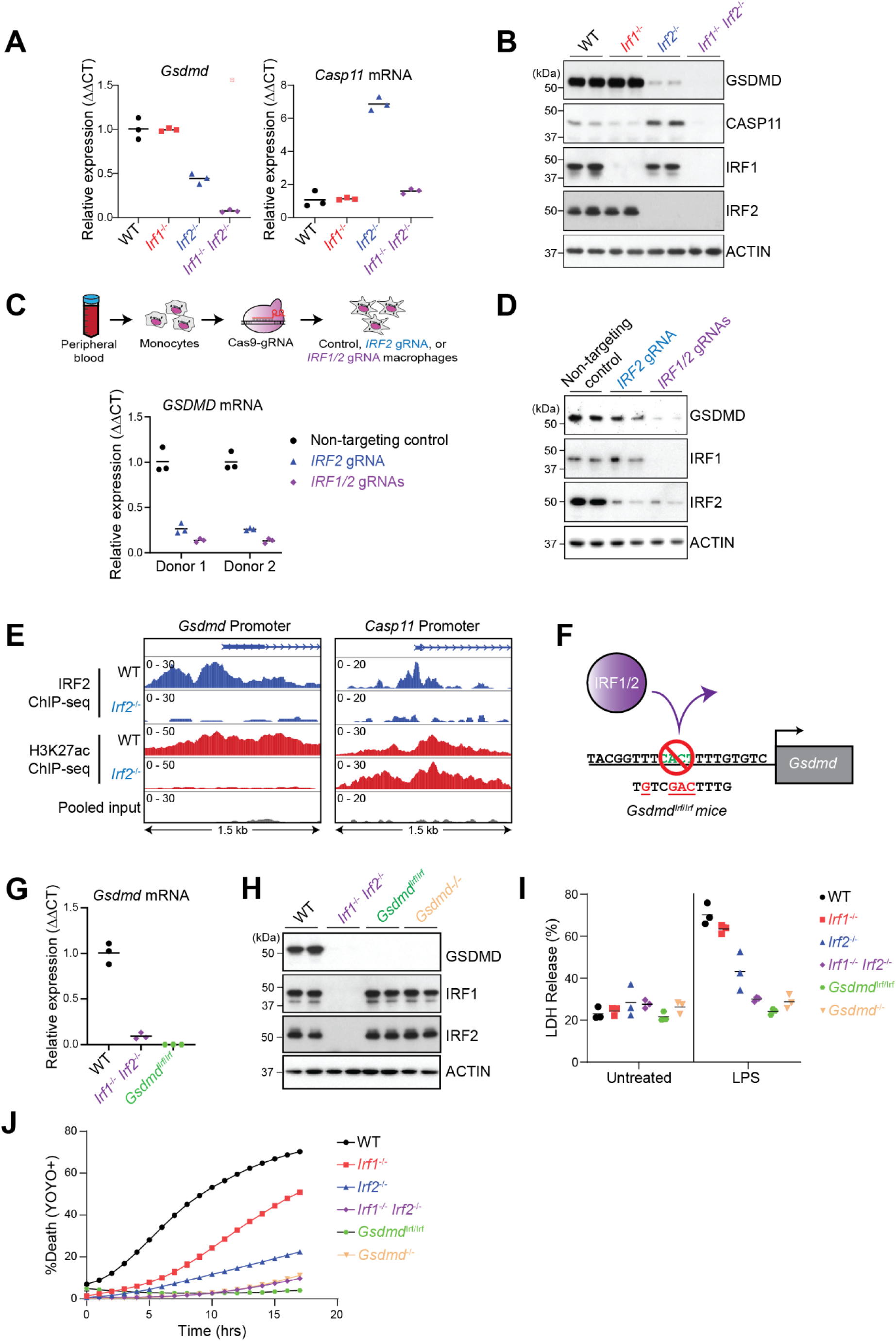
IRF1 transcribes *Gsdmd* in the absence of IRF2. (A and G) Relative *Gsdmd* and *Casp11* mRNA expression in BMDMs. Symbols, BMDMs from different mice (*n* = 3 per genotype). Lines indicate the mean. (B and H) Immunoblots of BMDMs (*n* = 2 per genotype). (C) Relative *GSDMD* mRNA expression in human peripheral blood monocyte (PBMC)-derived macrophages (*n* = 2 human blood donors) that were electroporated with the indicated gRNAs. Symbols, technical replicates. Results representative of 2 independent experiments. (D) Immunoblots of the PBMC-derived macrophages in (C). Results representative of 2 independent experiments. (E) IRF2 and H3K27ac ChIP-seq of mouse BMDMs at the *Gsdmd* and *Casp11* promoters. Numbers denote track scales (normalized read counts). Results representative of 2 biological replicates using cells from individual mice. (F) Sequence of the IRF binding site in the *Gsdmd* promoter of WT and *Gsdmd*^Irf/Irf^ mice. (I) LDH released from BMDMs that were primed with Pam3CSK4 for 4 h and then transfected with LPS for 18 hours. Symbols, BMDMs from 3 individual mice (*n* = 3 per genotype). Lines indicate the mean. (J) YOYO-1 uptake by BMDMs after priming with Pam3CSK4 for 4 hours, and then transfection with LPS. Datapoints are the mean. *n* = 3 per genotype.

**Figure S2.**
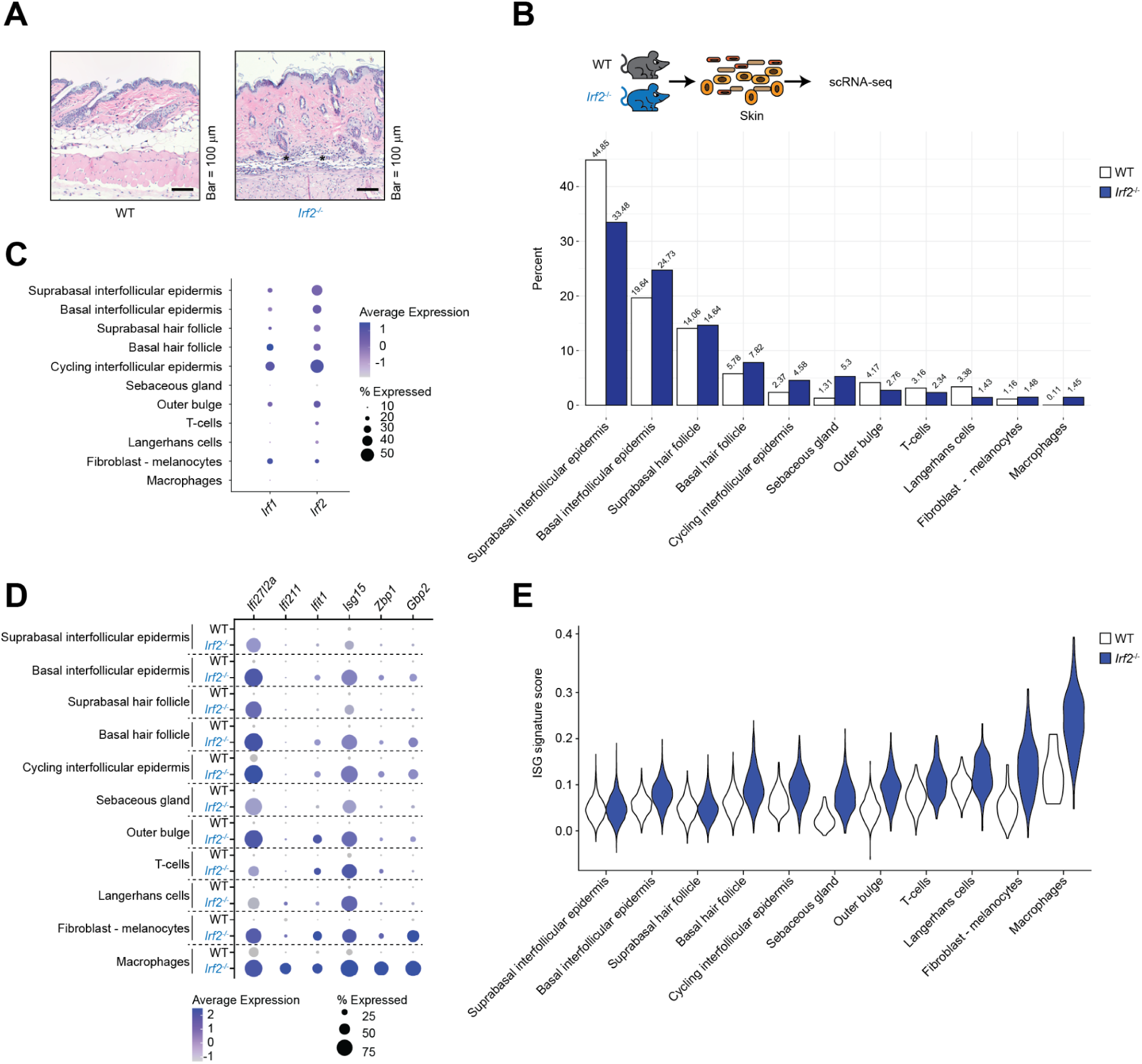
IRF2 suppresses ISG expression across all skin cell-types. Related to Figure 1. (A) Hematoxylin and eosin-stained skin sections of mice aged 12 weeks. Scale bar, 100 μm. (B) Bar graph shows the percentage of skin cells analyzed by scRNA-seq and clustered according to known skin markers ^20^. (C) Dot plot shows the mean level of *Irf1* and *Irf2* expression (dot intensity, blue scale) and the percentage of *Irf*-positive skin cells within each cell-type identified in (B). (D) Dot plot shows the mean level of expression of selected ISGs (dot intensity, blue scale) and the percentage of ISG-positive cells (dot size) within each population identified in (B). (E) ISG signature scores (genes defined in supplementary Table 1) for the cell-types in (B).

**Figure S3.**
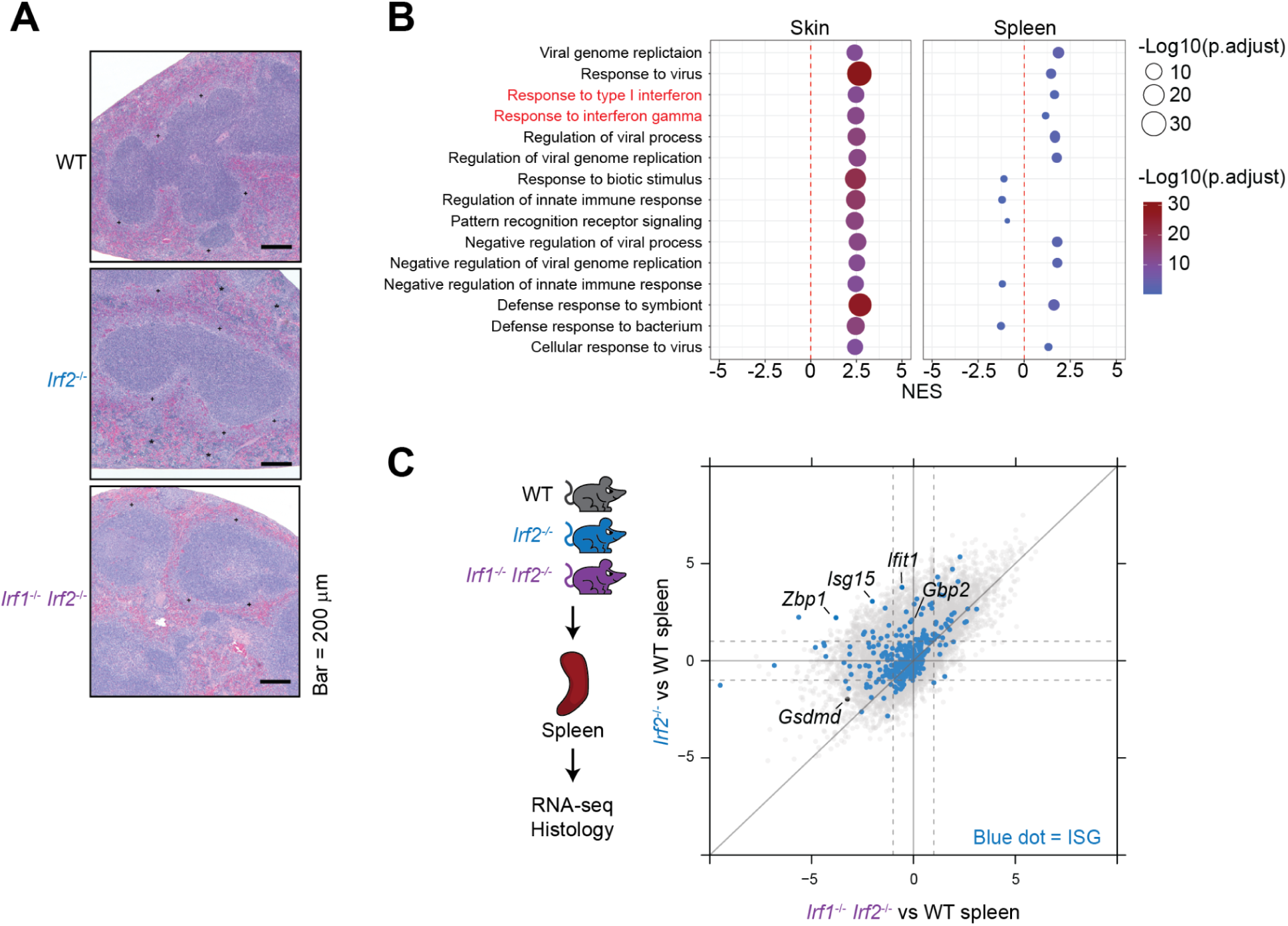
IRF1 drives elevated ISG expression in the absence of IRF2. Related to Figure 1. (A) Hematoxylin and eosin-stained spleen sections of mice aged 12 weeks. Results representative of 8 WT and 7 *Irf2^-/-^* mice. Asterisks denote zones of extramedullary haematopoiesis. Scale bar, 100 μm. (B) Enriched ontology terms for differentially expressed genes (log2 fold change >2 and *P* <0.05) in *Irf2*^-/-^ skin and spleen relative to WT. (C) Four-way plot of ISG expression (blue dots) in the spleen. Dashed lines mark log2 fold change cut-offs (<-1 or >1). ISGs elevated in both genotypes fall within the upper-right quadrant. ISGs elevated only in *Irf2*^-/-^ samples fall within the upper-left quadrant. *n* = 3 mice per genotype.

**Figure S4.**
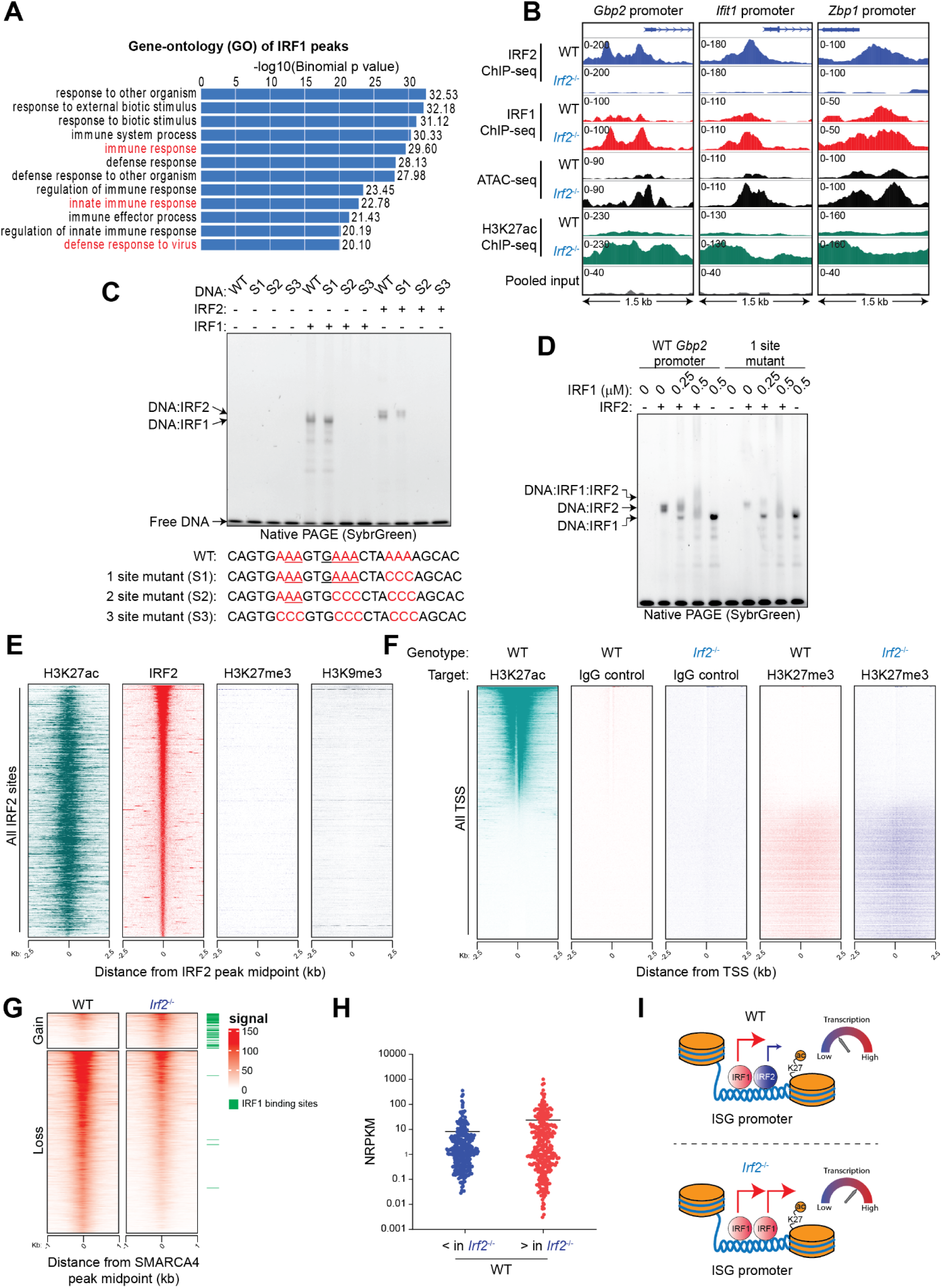
IRF2 limits gene expression at sites shared with IRF1. Related to Figure 1. (A) Enriched ontology terms for IRF1 ChIP-seq peaks found at promoters and enhancers (725 sites). (B) IRF2 ChIP-seq, IRF1 ChIP-seq, ATAC-seq, and H3K27ac ChIP-seq at ISG promoters in BMDMs. Numbers denote track scales (normalized read counts). Results representative of 2 biological replicates using cells from individual mice. (C) Gel shift assay of recombinant IRF1 and IRF2 (0.5 uM each) bound to a DNA sequence from the *Gbp2* promoter or its mutants. Gels were stained with SybrGreen to visualize the complexes. (D) Gel shift assay of recombinant IRF2 (0.5 uM) bound to the DNAs encoding WT or mutant *Gbp2* promoters in the presence of recombinant IRF1. Gels were stained with SybrGreen to visualize the complexes. (E) H3K27ac ChIP-seq, IRF2 ChIP-seq, H3K27me3 CUT&RUN, and H3K9me3 ChIP-seq in BMDMs. Plots show the distribution of normalized read counts in a 5,000 bp window (binned at 10 bps) around 4,822 IRF2 peaks. All rows are linked and sorted by total reads in a 1,000 bp region around IRF2 peaks genome-wide. (F) H3K27ac CHIP-seq and H3K27me3 CUT&RUN in WT and *Irf2*^-/-^ BMDMs. Plots show the distribution of normalized read counts in a 5,000 bp window (binned at 10 bps) around all TSS. All rows are linked and sorted by total reads in a 1,000 bp region around H3K27ac peaks in the WT sample. (G) Heatmap representation of SMARCA4 ChIP-seq at differentially bound sites. The signal is centered on SMARCA4 peaks in the WT sample, with a +/-1 Kb region shown. Green dashes represent sites shared by IRF1 and IRF2 (as shown in Figure 2E). Results representative of 2 biological replicates using cells from individual mice. (H) Expression of IRF2 activated or suppressed genes (log<2 or log>2, respectively, in *Irf2*^-/-^ vs WT skin) in WT skin. NRPKM, normalized reads per kilobase per million reads. Dots represent individual genes. *n* = 3 mice per genotype. Lines indicate the mean. (I) Model for IRF1 and IRF2 co-regulation of ISGs. IRF1 and IRF2 binding at the same genomic loci results in less transcription due to IRF2’s weaker activity relative to IRF1. In the absence of IRF2, more IRF1 binding results in increased ISG expression.

**Figure S5.**
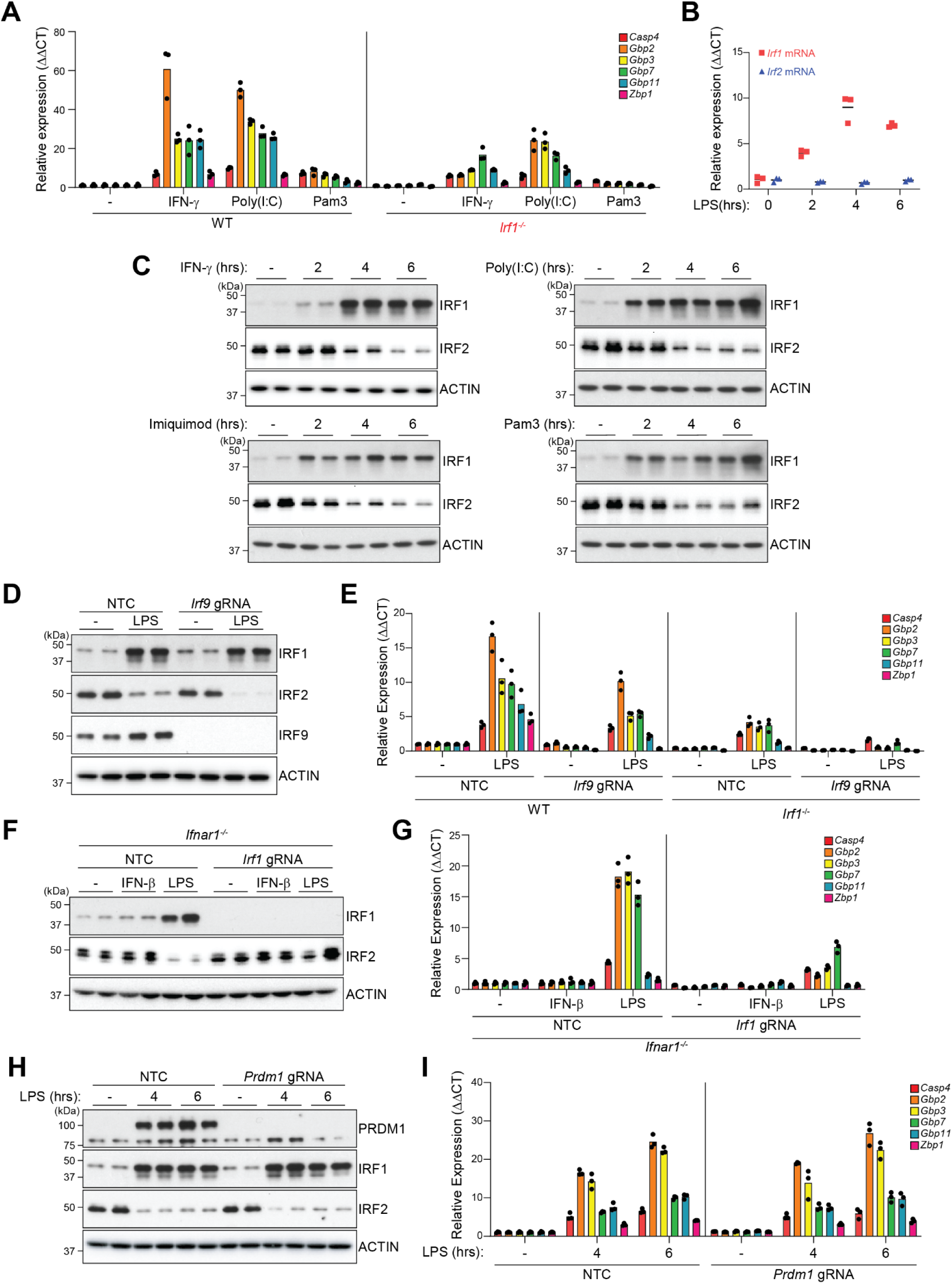
TLR signaling triggers proteasomal degradation of IRF2. Related to Figure 2. (A) RT-qPCR of ISGs in BMDMs treated with IFN-γ, Poly(I:C), or Pam3CSK4 for 4 hours. Mean expression levels of each gene are shown relative to those in untreated WT BMDMs. Dots, BMDMs from individual mice. Results representative of 3 independent experiments. (B) RT-qPCR of *Irf1* and *Irf2* in LPS-treated BMDMs. Mean expression levels of each gene are shown relative to those in untreated BMDMs. Dots, BMDMs from individual mice. Results representative of 3 independent experiments. (C) Immunoblots of BMDMs treated with IFN-γ, Poly(I:C), Imiquimod, or Pam3CSK4. Lanes represent BMDMs from individual mice (*n* = 2). (D) Immunoblots of BMDMs treated with LPS for 4 hours. Lanes represent BMDMs from individual mice (*n* = 2). Results representative of 3 independent experiments. (E) RT-qPCR of ISGs in BMDMs treated with LPS for 4 hours. Mean expression levels of each gene are shown relative to those in untreated WT BMDMs. Dots, BMDMs from individual mice. Results representative of 3 independent experiments. (F) Immunoblots of BMDMs treated with IFN-□ or LPS for 4 hours. Lanes represent BMDMs from individual mice (*n* = 2). Results representative of 3 independent experiments. (G) RT-qPCR of ISGs in BMDMs treated with IFN-□ or LPS for 4 hours. Mean expression levels of each gene are shown relative to those in untreated *Ifnar*^-/-^ BMDMs. Dots, BMDMs from individual mice. Results representative of 3 independent experiments. (I) Immunoblots of BMDMs treated with LPS. Lanes represent BMDMs from individual mice (*n* = 2). Results representative of 3 independent experiments. (H) RT-qPCR of ISGs in BMDMs treated with LPS. Mean expression levels of each gene are shown relative to those in untreated WT BMDMs. Dots, BMDMs from individual mice. Results representative of 3 independent experiments.

**Figure S6.**
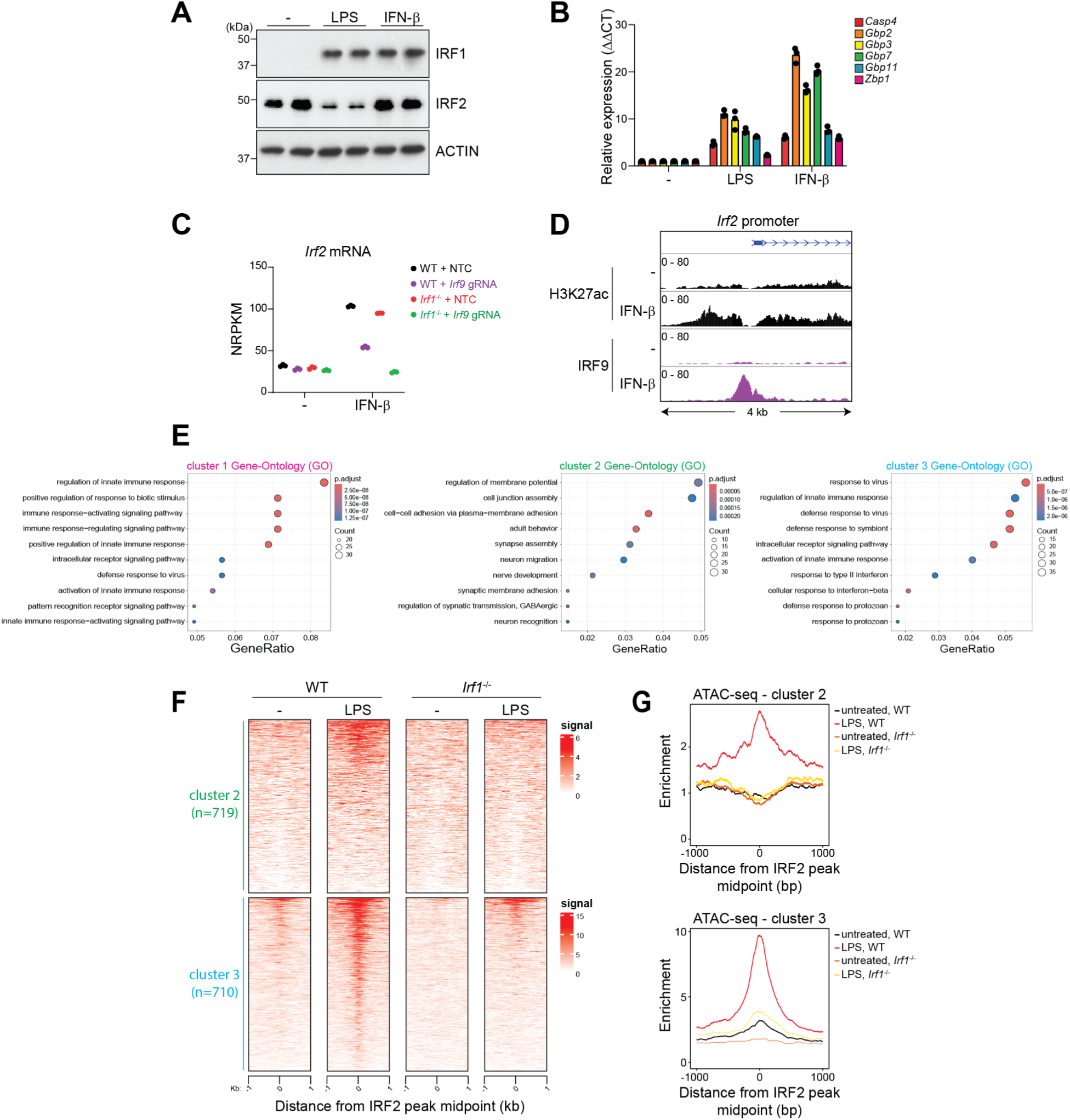
IRF9-mediated *Irf2* transcription masks IRF2 protein loss after IFN-□ stimulation. Related to Figure 4. (A) Immunoblots of BMDMs treated with IFN-□ or LPS for 4 hours. Lanes represent BMDMs from individual mice (*n* = 2). Results representative of 3 independent experiments. (B) RT-qPCR of ISGs in BMDMs treated with IFN-□ or LPS for 4 hours. Mean expression levels of each gene are shown relative to those in untreated BMDMs. Dots, BMDMs from individual mice. Results representative of 3 independent experiments. (C) *Irf2* expression in BMDMs treated with IFN-□ for 4 hours. NRPKM, normalized reads per kilobase per million reads. Symbols, BMDMs from individual mice (*n* = 3 per genotype). Lines indicate the mean. (D) IRF9 and H3K27ac ChIP-seq at the *Irf2* promoter in BMDMs treated with IFN-□ for 2 hours. Numbers denote track scales (normalized read counts). Results representative of 2 biological replicates using cells from individual mice. (E) Enriched ontology terms of sites in clusters 1, 2, and 3 (as shown in Figure 3D). (F) Heatmap representation of ATAC-seq at clusters 2 and 3 sites (as shown in Figure 3D) in BMDMs treated with LPS for 6 hours. The signal is centered on ATAC-seq peaks in the WT sample, with a +/-1 Kb region shown. Results representative of 2 biological replicates using cells from individual mice. (G) Density plots of ATAC-seq enrichment at cluster 2 and 3 in (F).

**Figure S7.**
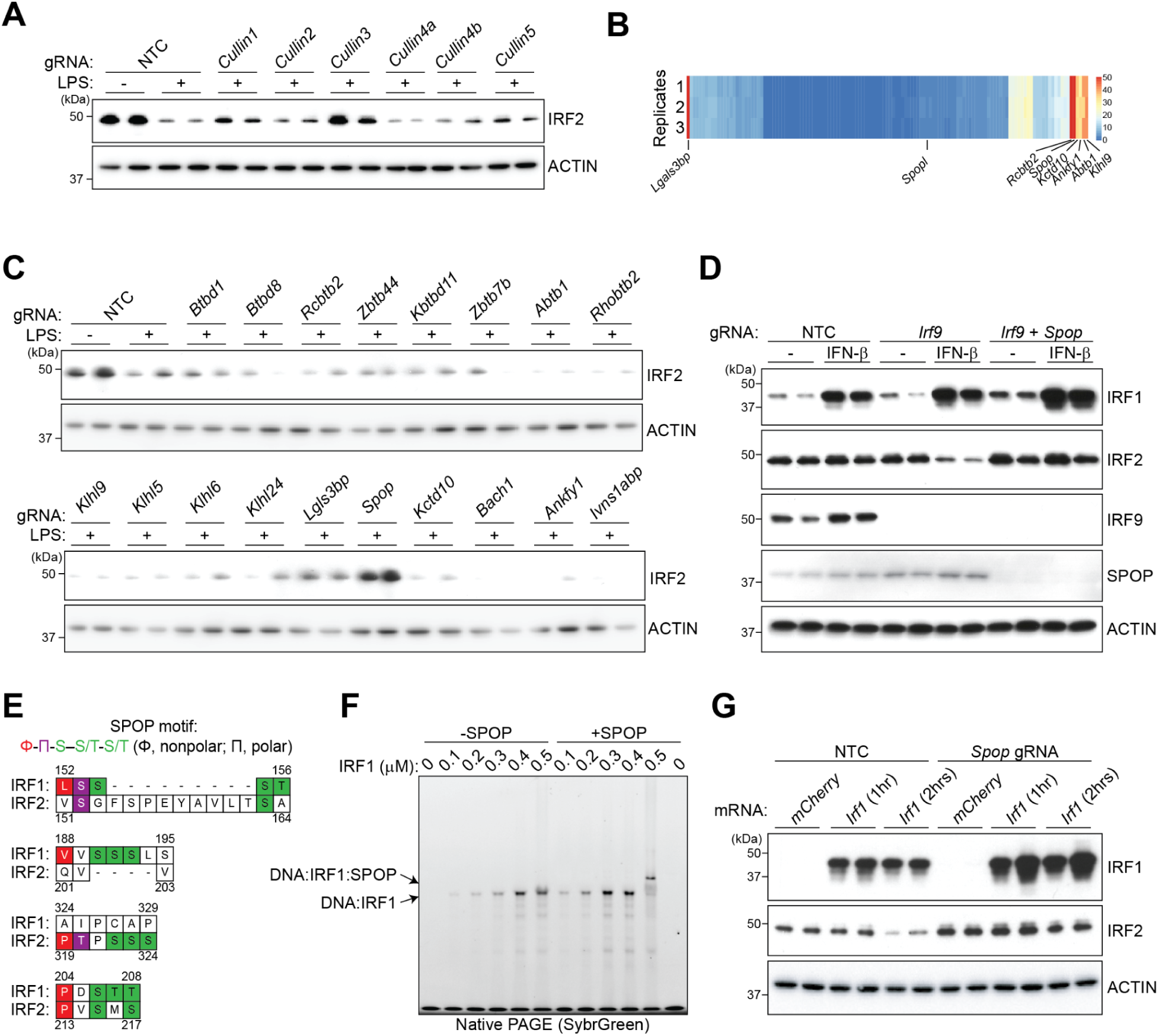
SPOP targets IRF2 for degradation. Related to Figure 4. (A and C) Immunoblots of BMDMs treated with LPS for 4 hours. Lanes represent BMDMs from individual mice (*n* = 2). Results representative of 3 independent experiments. (B) Heatmap showing the mRNA expression of Cullin 3 substrate adaptors in BMDMs. (D) Immunoblots of BMDMs treated with IFN-□ for 4 hours. Lanes represent BMDMs from individual mice (*n* = 2). Results representative of 3 independent experiments. (E) Diagram depicting the SPOP binding motifs found in mouse IRF1 and IRF2. (F) Gel shift assay of recombinant IRF1 bound to the DNAs encoding the *Gbp2* promoter in the presence of recombinant SPOP (2uM). Gels were stained with SybrGreen to visualize the complexes. (G) Immunoblots of BMDMs electroporated with *Irf1* or *mCherry* mRNAs. Lanes represent BMDMs from individual mice (*n* = 2). Results representative of 3 independent experiments.

**Figure S8.**
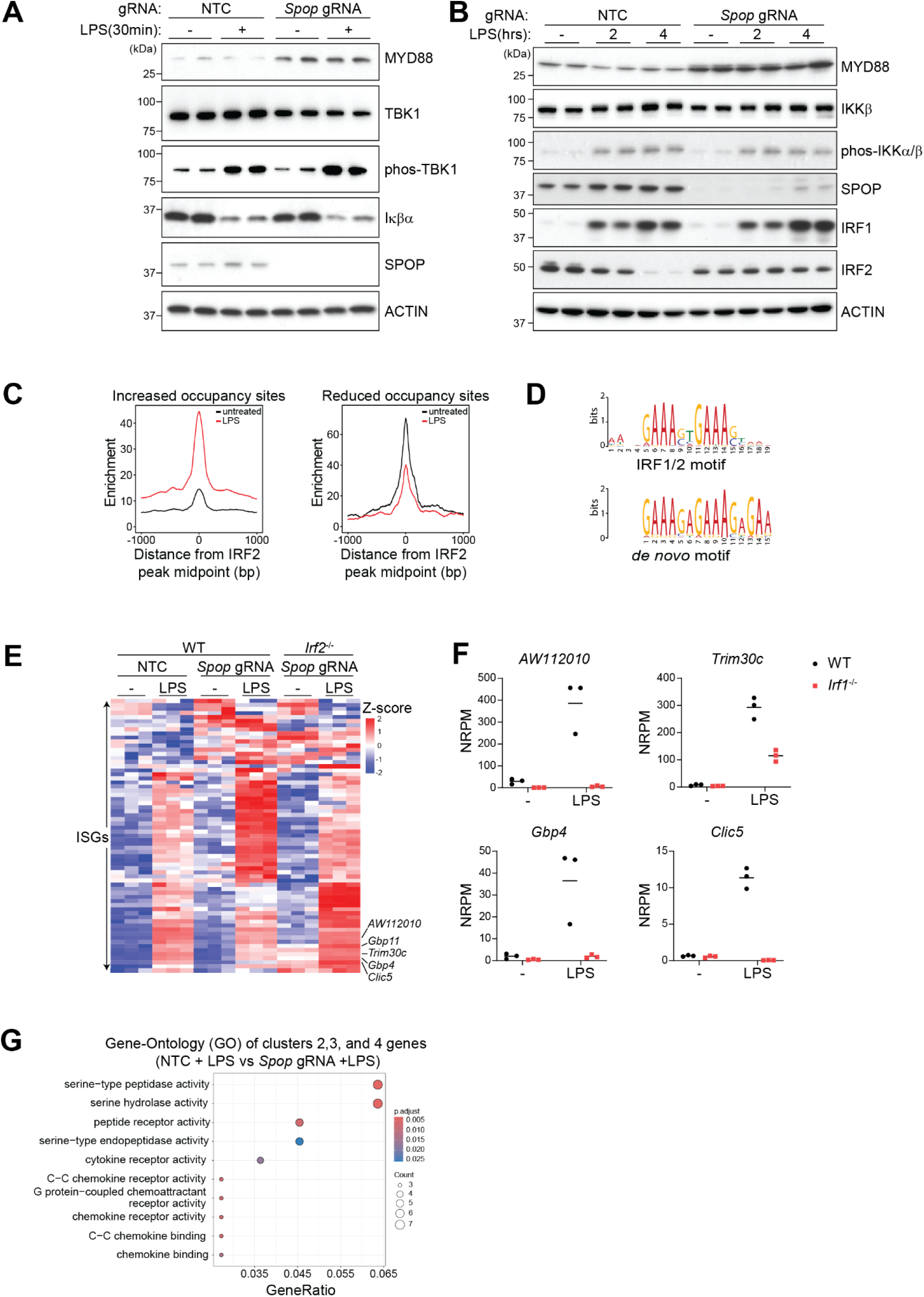
IRF2 drives aberrant gene transcription in *Spop*-deficient BMDMs. Related to Figure 5. (A) Immunoblots of BMDMs treated with LPS for 30 minutes. Lanes represent BMDMs from individual mice (*n* = 2). Results representative of 3 independent experiments. (B) Immunoblots of BMDMs treated with LPS. Lanes represent BMDMs from individual mice (*n* = 2). Results representative of 3 independent experiments. (C) Density plots of differentially bound IRF2 sites in Figure 5A. (D) *de novo* motif analysis at sites of increased IRF2 occupancy in Figure 5A. (E) Heat-map shows gene expression (Z-score) for 67 differentially expressed ISGs (adjusted *P* <0.05, log2 fold change <-1 or >1) in BMDMs treated with LPS for 6 hrs (*n* = 3 mice per genotype). (F) Expression of selected ISGs in BMDMs treated with LPS for 6 hours. NRPKM, normalized reads per kilobase per million reads. Symbols, BMDMs from individual mice (*n* = 3 per genotype). Lines indicate the mean. (F) Enriched ontology terms for differentially upregulated genes in clusters 2, 3, and 4 of Figure 5E.

**Figure S9.**
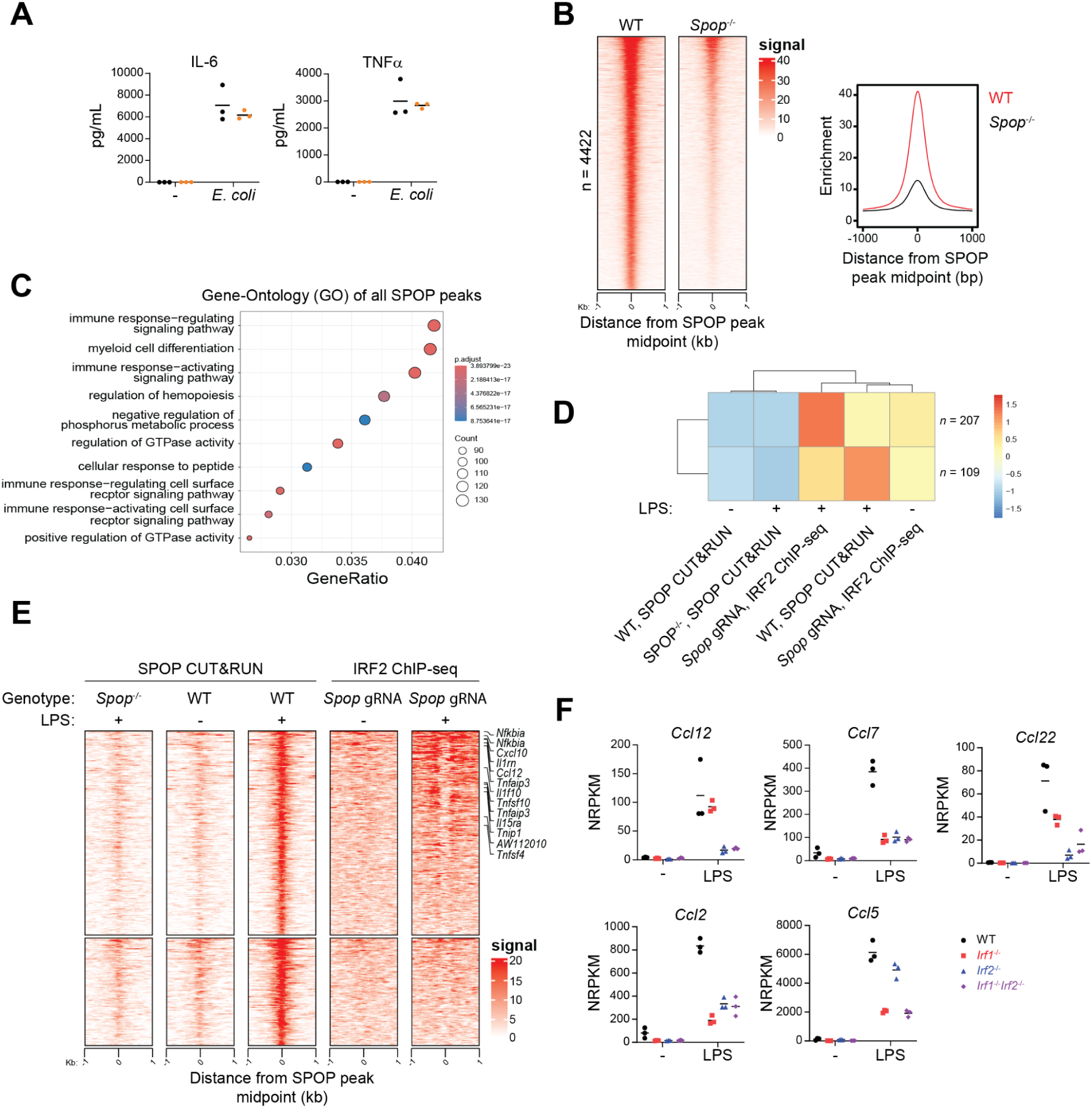
SPOP targets IRF2 at cytokine loci. Related to Figure 6. (A) IL-6 and TNFα secreted from BMDMs in Figure 6A. (B) Left: Heatmap representation of SPOP CUT&RUN peaks in BMDMs. The signal is centered on the SPOP peaks in WT BMDMs, with a +/-1 Kb region shown. Results representative of 2 biological replicates using cells from individual mice. Right: density plots of SPOP peaks. (C) Enriched ontology terms for SPOP peaks in (B). (D) K-means clustering of SPOP CUT&RUN and IRF2 ChIP-seq datasets at increased occupancy SPOP peaks from Figure 6C. (E) Heatmap representation of clusters in (D). The signal is centered on SPOP peaks in WT cells stimulated with LPS, with a +/-1 Kb region shown. Results representative of 2 biological replicates using cells from individual mice. (F) Cytokine mRNA expression in BMDMs treated with LPS for 6 hours. NRPKM, normalized reads per kilobase per million reads. Symbols, BMDMs from individual mice (*n* = 3 per genotype). Lines indicate the mean.

## Methods

### Mice

All mouse studies complied with relevant ethics regulations and were approved by the Genentech Institutional Animal Care and Use Committee in an Association for Assessment and Accreditation of Laboratory Animal Care (AAALAC)-accredited facility in accordance with the Guide for the Care and Use of Laboratory Animals and applicable laws and regulations. *Irf2*^-/-^ and *Gsdmd*^-/-^ mice with a C57BL/6N background were described previously ^18, 40^. The *Gsdmd^Irf/Irf^* mice were generated using C57BL/6N C2 ES cells. A targeting construct with a FRT-Neo-FRT selection cassette located 136 bp upstream of the 4 substitutions (ttcact => **g**tc**gac**, corresponding to GRCm39 chr15:75,734,179-75,734,184) was used to modify the *Gsdmd* promoter region immediately upstream of exon 1. After successful targeting, the Neo cassette was excised in ES cells using FLP, leaving a single FRT site. We used established CRISPR methods and C57BL/6N zygotes to obtain *Irf1*^-/-^ mice lacking a 2299 bp region that contains *Irf1* exons 3-7 (GRCm3 chr11:53,663,345-53,665,643). The sgRNAs used were 5’-GCUACCGAGCAUUGGAACAU-3’ and 5’-AGCCACCUUUGACCAAGUUG-3’. To obtain the Spop conditional knockout allele, two loxP sites were inserted flanking a 1,280 bp region containing exons 4-5, corresponding to chr11:95,364,795-95,366,074 (GRCm39/mm39). The following sgRNA targets were used: 5’-CAGGTTAATGATTTGGCAAT-3’ and 5’-CCTTTCGATTAGATCTTGGG-3’. The two sgRNAs were co-electroporated with two Alt-R-HDR loxP donor oligos (Integrated DNA Technologies) and G0 animals were screened using NGS and in vitro cre recombination to identify cis-targeted founders. N1 mice were screened using NGS, in vitro cre recombination and dPCR to identify animals carrying the correctly targeted Spop conditional knockout allele. Potential off-targets for gRNA were interrogated using rhAmpSeq analysis, and N1 heterozygous mice without off-targets were used to establish the line. All mouse alleles were maintained on a C57BL/6N genetic background.

### Cell culture

Bone marrow cells were differentiated into macrophages in DMEM supplemented with 10% heat-inactivated foetal bovine serum (FBS), 2 mM glutamine, 1× non-essential amino acids solution, 100 U/ml penicillin, 100 μg/ml streptomycin, and 20% (v/v) L929-conditioned medium at 37°C with 5% CO2. BMDMs were collected on day 5 and seeded for experiments overnight. HEK293T cells (ATCC CRL-3216, mycoplasma tested but not authenticated) were cultured in DMEM supplemented with 10% heat-inactivated FBS, 2 mM glutamine, 1× non-essential amino acids solution, 100 U/ml penicillin and 100 μg/ml streptomycin.

Peripheral blood and PBMCs were collected from healthy donors participating in the Genentech blood donor program after written, informed consent from the Western Institutional Review Board. PBMCs were isolated from donor blood using Sepmate tubes (StemCell Technologies, 85450). Red blood cells were lysed using ACK lysing buffer (ThermoFisher, A1049201). Monocytes were isolated from PBMCs using the EasySep Human Monocyte isolation kit (Stem Cell Technologies, 19359).

### CRISPR-mediated gene editing

CRISPR-mediated gene editing of freshly isolated human monocytes was performed as described ^41^. Guide RNAs targeting *IRF1* (Hs.Cas9.IRF1.1.AB) and *IRF2* (Hs.Cas9.IRF2.1.AB) were from IDT. The sequence of the non-targeting control gRNA was CGTTAATCGCGTATAATACG. gRNAs were complexed with Cas9 (IDT, 1081059) at room temperature for 10 min. Monocytes (1.5×10^6^) were electroporated with the gRNA-Cas9 complex using the P3 Primary Cell 4D-Nucleofector Kit (Lonza, V4XP-3032) and a Lonza 4D-nucleofector system (Lonza, AAF-1002B and AAF-1002X; CM-137 settings). Monocytes were seeded into 24-well plates (5×10^5^/well) containing 1 ml macrophage medium (DMEM supplemented with 10% heat-inactivated FBS, 2 mM glutamine, 1× non-essential amino acids solution, 100 U/ml penicillin, 100 μg/ml streptomycin, and 100 ng/ml human recombinant M-CSF [Peprotech, 300-25]) and differentiated into macrophages for 5 days.

BMDMs were differentiated for 5 days and CRISPR-edited using the same protocol. gRNAs targeting two sites in *Irf1*, *Irf9*, *Spop*, and *Prdm1* were purchased from IDT (Mm.Cas9.IRF1.1.AA, Mm.Cas9.IRF1.1.AB, Mm.Cas9.IRF9.1.AA, Mm.Cas9.IRF9.1.AB, Mm.Cas9.SPOP.1.AA, Mm.Cas9.SPOP.1.AB, Mm.Cas9.PRDM1.1.AA, Mm.Cas9.PRDM1.1.AB). Electroporated BMDMs were seeded into 24-well plates (5×10^5^/well) containing 1 mL BMDM media and stimulated after 3 additional days.

### Stimulations

BMDMs were stimulated in Opti-MEM (ThermoFisher, 31985062) with 100 ng/ml ultrapure LPS (InvivoGen, O111:B4), 5 ng/ml mouse IFN-□ (R&D Systems, 8234-MB-010/CF), 5 ng/ml mouse IFN-γ (BioLegend, 575302), 1 μg/ml Pam3CSK4 (InvivoGen, tlrl-pms), 1 μg/ml Imiquimod (InvivoGen, tlrl-imqs-1), 50 μg/ml pHrodo Green *S*. *aureus* bioparticles (ThermoFisher, P35367), 50 μg/ml pHrodo Green *E*. *coli* Bioparticles (ThermoFisher, P35366), 5 μM MG-132 (Selleckchem, S2619), 0.1 μM MLN4924 (ChemieTek, CT-M4924), 0.5 or 1 μM Baricitinib (MedChemExpress, HY-15315), 0.5 or 1 μM Ruxolitinib (InvivoGen, tlrl-rux-3), and 0.5 μg/mL Actinomycin D (Sigma, A1410).

For ectopic expression of IRF1, *Irf1*^-/-^ BMDMs (5×10^5^) were electroporated with 500ng *Irf1* (GenScript, NM_008390.2) or mCherry mRNA (GenScript, SC2325), then seeded in 24-well plates in BMDM media.

BMDMs in 96-well format (1×10^5^/well) were primed with 1 μg/ml Pam3CSK4 (InvivoGen, tlrl-pms) for 4 h, and then transfected with 5 μg/ml LPS and 0.25% (v/v) FuGENE HD (Promega, E2311) in Opti-MEM. To monitor cell survival, the media was supplemented with 200 nM YOYO-1 (ThermoFisher, Y3601) and images were collected in an IncuCyte ZOOM (Essen BioScience) at ×10 magnification. Nuclear-ID Red DNA stain (Enzo Life Sciences, ENZ-52406) was added at a 500-fold dilution at the last time point. IncuCyte software was used to enumerate YOYO-1+ and Nuclear-ID+ cells. Percentage cell death was calculated as the number of YOYO+ cells divided by the number of Nuclear-ID+ cells.

### Immunoblotting

BMDMs (5×10^5^) were lysed in 45 μl of SDS buffer (1% SDS, 150 mM NaCl, 50 mM tris pH 7.5, 1 mM EDTA) and incubated at 90°C for 10 min. Protein concentration was determined using the Pierce™ BCA Protein Assay Kit (ThermoFisher, 23225). Antibodies recognized ACTIN (CST, 13E5), IRF1 (CST, D5E4), IRF2 (Abcam, ab124744), IRF9 (Millipore, 6F1-H5), GSDMD (Genentech, 17G2G9) ^42^, caspase-11 (Novus, NB120-10454), PRDM1 (Abcam, ab241568), phospho-STAT1 (CST, 9167S), STAT1 (CST, 9172S), SPOP (Proteintech, 16750-1-AP), MYD88 (CST, 4283S), phospho-TBK1 (CST, 5483S), TBK1 (CST, 38066S), IκBα (CST, 4814S), IKKβ (CST, 8943S), and phospho-IKKα/β (CST, 2078S).

### SPOP immunoprecipitation

BMDMs from 2-3 mice were pooled per IP reaction. The cytosolic fraction was removed with mild lysis buffer (0.2% NP-40, 25 mM TRIS pH 7.5, 150 mM NaCl, 2 mM MgCl_2_) containing EDTA-free protease inhibitor cocktail (ThermoFisher, 78425), 25 U/mL benzonase (Millipore Sigma, 70664-3), and 10 μM MG-132 (Selleckchem, S2619). Nuclear fractions were lysed with RIPA buffer (ThermoFisher, 89900) containing EDTA-free protease inhibitor cocktail (ThermoFisher, 78425), 25 U/mL benzonase (Millipore Sigma, 70664-3), and 10 μM MG-132 (Selleckchem, S2619). SPOP complexes were captured from BMDM lysates with 1 μg SPOP antibody (Genentech, clone F5) and protein A/G magnetic beads (ThermoFisher, 88802). Beads were extensively with RIPA buffer and eluted in soft elution buffer (0.2% SDS, 0.1% Tween 20) at 65°C for 3 minutes.

### Protein purification

N-terminally myc-DDK-tagged full-length mouse IRF2 (OriGene, MR205275) was expressed in HEK293T cells for 48 hours. Cells were lyzed in a buffer containing 0.2% NP-40, 10% glycerol, 300 mM NaCl, 50 mM Na_3_PO_4_, 10 mM imidazole, 2 mM MgCl_2_, EDTA-free protease inhibitor cocktail (ThermoFisher, 78425) and 25 U/mL benzonase (Millipore Sigma, 70664-3). The protein was purified by a combination of FLAG immunoprecipitation with anti-FLAG® M2 magnetic beads (Millipore Sigma, M8823) and size exclusion chromatography (SEC) in buffer containing 20 mM HEPES pH 7.5, 150 mM NaCl and 2 mM DTT.

N-terminally MBP-tagged mouse SPOP (aa28-329) was cloned into the pAcSg2 vector expressed in Tni insect cells for 48hrs. Cells were lysed using a Dounce homogenizer in a buffer containing 40mM HEPES pH 7.5, 300mM NaCl, 0.1mM TCEP, and protease inhibitors (ThermoFisher, 78425). The protein was purified by a combination of amylose resin pull-down (New England Biolabs, E8021S), TEV cleavage (GenScript, Z03030), and size exclusion chromatography (SEC) in buffer containing 30 mM HEPES pH 7.5, 150 mM NaCl and 0.1 mM TCEP.

N-terminally FLAG-tagged full-length mouse IRF1 was cloned into the pAcSg2 vector and expressed in Tni insect cells for 48hrs. Cells were lysed using a Dounce homogenizer in a buffer containing 40mM HEPES pH 7.5, 150mM NaCl, 1 mM MgCl_2_, 2 μg//mL leupeptin, 160 μg/mL benzamidine, and 0.1mM TCEP. The protein was purified by a combination of FLAG immunoprecipitation with anti-FLAG® M2 affinity gel (Millipore Sigma, A2220) and size exclusion chromatography (SEC) in buffer containing 30 mM HEPES pH 7.5, 150 mM NaCl, 10% glycerol, and 0.1 mM TCEP.

### Native gel shift assays

Double-stranded DNA encoding the IRF binding site in the mouse *Gbp2* promoter (2 ng/μL) was incubated with recombinant IRF1, IRF2, or SPOP (at the indicated concentrations) in EMSA buffer (20 mM HEPES, 50 mM NaCl, 1.5 mM MgCl_2_, 5 mM DTT) for 10 minutes at room temperature. Complexes were run on 4–12% Tris-Glycine gels (ThermoFisher, XP04122BOX) with Tris-Glycine native running buffer (ThermoFisher, LC2672). Gels were stained with SYBR Green (ThermoFisher, S7563) and imaged using the Gel Doc EZ Imager (Bio-Rad).

### Histology and Immunohistochemistry

Tissues were formalin-fixed, paraffin-embedded, and stained with haematoxylin and eosin for histologic evaluation. Skin lesions were scored in a blind and randomised manner, and included assessments of inflammation, epidermal hyperplasia, and dermal edema. Inflammation was scored as: (1) ≤1 inflammatory focus per skin section, (2) multifocal to locally extensive dermal and/or subcutaneous inflammation, or (3) extensive inflammation with disruption of tissue architecture. Epidermal hyperplasia was scored as: (1) focal epidermal thickening of less than 3 follicles in length, (2) segmental epidermal thickening (3 or more cell layers) of ≤50% of the skin section, or (3) thickening of >50% of the epidermis. Dermal edema was scored as: (1) localised subepidermal edema, (2) multifocal to segmental dermal edema, or (3) diffuse edema with distortion of the tissue architecture. Inflammation, epidermal hyperplasia, and edema scores were summed to generate a final score for each mouse.

ZBP1 immunohistochemistry on formalin-fixed, paraffin-embedded tissue sections was performed on a Dako autostainer (Agilent) with GN58.3 rat anti-mouse ZBP1 antibody (Genentech). Antigen retrieval was performed with EDTA pH 8 (Abcam) and non-specific labelling was blocked with Sytek biotin block and hydrogen peroxide. ZBP1 was labelled with 7.5 µg/ml anti-ZBP1 antibody for 1 h at room temperature. Bound antibody was detected with biotinylated donkey anti-rat IgG (H+L) antibody (Jackson ImmunoResearch) and Vectastain Elite ABC-HRP (Vector Laboratories) with diaminobenzidine as the chromogen. Slides were counterstained with Mayer’s hematoxylin. A naïve rat IgG2b antibody (Pharmingen) was used in place of the primary antibody as a negative control, and lacked significant labelling. Additional controls included WT and *Zbp1^-/-^* mouse embryos, which displayed expected differential labelling.

### RNA isolation and qPCR

Total RNA was extracted using a RNeasy Mini Kit (Qiagen, 74134). Skin and spleen samples were homogenised in the RLT buffer with ceramic beads (OMNI International, 19-628) and a Bead Ruptor 12 homogenizer (OMNI International). Total RNA was quantified in a NanoDrop 8000 (ThermoFisher). 250 ng of total RNA was used for reverse transcription using a High-Capacity cDNA Reverse Transcription Kit (ThermoFisher, 4368814). cDNA was diluted 10-fold in water and used for quantitative RT-PCR with TaqMan™ Universal PCR Master Mix (ThermoFisher, 4304437) and TaqMan probes [*Actb* (Mm02619580_g1), *Gsdmd* (Mm00509958_m1), *Gbp2* (Mm00494576_g1), *Gbp3* (Mm00497606_m1), *Gbp7* (Mm00523797_m1), *Gbp11* (Mm01613158_m1), *Isg15* (Mm01705338_s1), *Zbp1* (Mm01247052_m1), *Casp11* (Mm00432304_m1), *Tlr3* (Mm01207404_m1), *Irf1* (Mm01288580_m1), *Ccl7* (Mm00443113_m1), *Ccl8* (Mm01297183_m1), *Ccl2* (Mm00441242_m1), *Ccl12* (Mm01617100_m1), *Ccl22* (Mm00436439_m1), *ACTB* (Hs01060665_g1), and *GSDMD* (Hs00986745_g1)]. *Actb* or *ACTB* served as internal control genes. Target gene expression was normalised against the WT or untreated controls.

### RNA-seq

Total RNA was quantified with the Qubit RNA HS Assay Kit (ThermoFisher, Q32852) and assessed using the RNA ScreenTape on TapeStation 4200 (Agilent Technologies). Sequencing libraries were generated using the Truseq Stranded mRNA kit (Illumina, 20020594) with an input of 100-1,000 ng of total RNA (for BMDM experiments), or the SMARTer Stranded Total RNA-Seq Kit v2 - Pico Input Mammalian (Takara, 634411) with 1-10 ng of total RNA (for skin or spleen experiments). Libraries were quantified with the Qubit dsDNA HS Assay Kit (ThermoFisher, Q32851) and the average library size was determined using the D1000 ScreenTape on TapeStation 4200 (Agilent Technologies). Libraries were pooled and sequenced on NovaSeq 6000 (Illumina) to generate 30 million single-end 50-base pair reads for each sample. GSNAP (version 2013-11-10) ^43^ was used to align raw FASTQ reads to the reference genome (mouse GRCm38/mm10 or human GRCh38) with following parameters “-M 2 -n 10 -B 2 -i 1 -N 1 -w 200000 -E 1 --pairmax-rna=200000 --clip-overlap”. Only uniquely mapped reads were used for downstream analyses. Limma (version 3.54.1) ^44^ R package was used to perform differential expression analysis. Heatmaps were plotted using R package pheatmap (version 1.0.12). Gene set enrichment analysis was done using R package clusterProfiler (version 4.6.0) ^45^. *P* values were calculated using a moderated version of t-test. Resulting *P* values were adjusted for multiple comparisons by controlling the false discovery rate (FDR) using the Benjamini-Hochberg method.

### scRNA-seq

Skin cells were isolated as described ^46^ with slight modifications. Mouse skin taken from the chest was digested with 1000 U/ml collagenase (Gibco, 17104019) and 20 U/ml DNase I (Sigma-Alrich, DN25) in cell dissociation buffer (ThermoFisher, 13150016) for 10 min at 37°C. Skin samples were digested two additional times, replenishing the digestion solution after spinning samples down. Tissue was mechanically disrupted by pipetting after each digestion. Single cell suspensions were filtered through 70 µm cell strainers and processed using the 10x Genomics Chromium Single Cell 3′ V3.1 Reagent Kit (PN-1000121). For each sample, we targeted 10,000 viable cells. GEMs were generated, cells were lysed, and mRNAs were barcoded and underwent reverse transcription to produce full-length cDNAs. cDNAs were amplified with 12 PCR cycles. Gene expression libraries were sequenced at a depth of 20,000 reads/cell on an Illumina NovaSeq6000.

10X 3’ single cell RNA-sequencing data was processed with cellranger (version 7.1.0) ^47^. A gene by barcode matrix was generated by mapping reads to the mouse reference genome (GRCm38). Seurat (version 4.0.4) ^48^ was used for downstream analyses on cells with between 200 and 4,000 detected genes (9,395 WT cells and 11,486 *Irf2*^-/-^cells). After log normalisation, dimensionality reduction was performed using principal component analysis (PCA). Principal components (PCs) were then ranked by the amount of variance explained, and after 25 PCs the explained variance plateaued. Clusters were then identified using the first 25 PCs with resolution of 0.5. The same PCs were used to generate UMAP projections. WT and *Irf2*^-/-^ samples were integrated by identifying anchor points between the two datasets. Cell annotation was performed using singleR using the reference atlas from Joost S. et al ^20^. The ISG signature was generated using the AddModuleScore function in Seurat and a list of known ISGs ^49^.

### ChIP-seq

H3K27ac, IRF2, and IRF1 ChIP was performed by Active Motif (Carlsbad, CA, USA). BMDMs were fixed with 1% formaldehyde for 15 min and quenched with 0.125 M glycine. Chromatin was isolated by the addition of a lysis buffer, followed by disruption with a Dounce homogenizer (Active Motif, 40401). Lysates were sonicated and the DNA sheared to an average length of 300-500 bp. Genomic DNA (input) was prepared by treating aliquots of chromatin with RNase (Sigma-Alrich, R5250), proteinase K (Sigma-Alrich, P6556) and heat (overnight, 65°C) for de-crosslinking, followed by SPRI beads clean up (Beckman Coulter, B23318) and quantitation by CLARIOstar microplate reader. Extrapolation to the original chromatin volume allowed determination of the total chromatin yield. For ChIP, chromatin (30 μg) was precleared with protein A-agarose (Invitrogen, 15918014) and then protein-DNA complexes were captured with 5 μg of antibody against IRF2 (Abcam, ab124744), IRF1 (CST, D5E4), or H3K27Ac (Active Motif, 39133), plus protein A-agarose. Complexes were washed, eluted with SDS buffer, and treated with RNase and proteinase K. Crosslinks were reversed by incubation overnight at 65°C, and ChIP DNA was purified by phenol-chloroform extraction and ethanol precipitation.

SMARCA4 ChIP was performed as previously described with the following modifications ^50^. BMDMs were crosslinked with 50 mM DSG (disuccinimidyl glutarate, ProteoChem, C1104) for 30 min and then with 1% formaldehyde for 10 min. Formaldehyde was quenched by the addition of glycine. Nuclei were prepared with ChIP lysis buffer (1% Triton x-100, 0.1% SDS, 150 mM NaCl, 1mM EDTA, and 20 mM Tris, pH 8.0) and sheared with a Covaris sonicator (Fill level – 10, Duty Cycle – 15, PIP – 350, Cycles/Burst – 200, Time – 8 min). Sheared chromatin was immunoprecipitated overnight with SMARCA4 antibody (Abcam, ab110641). Antibody chromatin complexes were captured with Protein A-magnetic beads and washed twice in buffer I (1% Triton, 0.1% SDS, 150 mM NaCl, 1 mM EDTA, 20 mM Tris pH 8.0, and 0.1% NaDOC), twice in buffer II (1% Triton, 0.1% SDS, 500 mM NaCl, 1 mM EDTA, 20 mM Tris, pH 8.0, and 0.1% NaDOC), twice in buffer III (0.25 M LiCl, 0.5% NP-40, 1mM EDTA, 20 mM Tris, pH 8.0, 0.5% NaDOC), and once in TE buffer (10 mM EDTA and 200 mM Tris, pH 8.0). DNA was eluted by vigorous shaking for 20 min in 100 mM NaHCO3, 1% SDS. DNA was de-crosslinked overnight at 65°C and purified with a MinElute PCR purification kit (Qiagen).

Illumina sequencing libraries were prepared from the ChIP and input DNAs by end-polishing, dA-addition, and adaptor ligation on an automated system (Apollo 342, Wafergen Biosystems/Takara). After a final PCR amplification step, the resulting DNA libraries were quantified and sequenced on Illumina’s NextSeq 6000 (75 nt reads, single end). The raw FASTQ files were aligned to the mouse reference genome (GRCm38) using BWA (version 0.7.12-r1039) ^51^. Reads that mapped uniquely to the genome and aligned with no more than 2 mismatches were used for downstream analyses. MACS2 (version 2.1.0) ^52^ was used to call the peaks relative to the pooled input. Differential peak analysis was done using DiffBind (version 3.16) R package ^53^. GREAT (version 4.0.4) ^54,55^ web application was used to perform gene ontology analysis. ChIPSeeker ^56^ was used to annotate peaks. SoGGI ^57^ and Profileplyr ^58^ were used to generate heatmaps and K-means clustering. MEME ^59^ was used for motif analysis. WashU epigenome browser ^60^ was used to plot the locus specific ChIP-seq read enrichment. ChIP-seq data for IRF9 were downloaded from GSE115435.

### ATAC-seq

Libraries were prepared with the ATAC-seq kit from Active Motif (53150). Libraries were pooled and sequenced on NovaSeq 6000 (Illumina) to generate 30 million single-end 50-base pair reads for each sample. ATAC-seq data was processed using the ENCODE ATAC-seq pipeline with minor changes. In brief, FASTQ adapter trimming was performed using cutadapt (version 1.9.1). The trimmed FASTQs were then aligned using the Bowtie aligner (version 2.2.6) ^61^ and only properly paired reads were used for downstream analyses. Picard (version 1.126) was used to mark duplicate reads. Peaks were called using MACS2 (version 2.1.0) ^52^ and were filtered to remove blacklisted regions. DiffBind (version 3.16) ^53^ R package was used to perform differential peak analysis. GREAT (version 4.0.4) ^54,55^ web application was used to perform gene ontology analysis. ChIPSeeker ^56^ was used to annotate peaks. SoGGI ^57^ and Profileplyr ^58^ were used to generate heatmaps and K-means clustering. WashU epigenome browser ^60^ was used to plot the locus specific ATAC-seq read enrichment.

### CUT&RUN

Reactions and Libraries were prepared with the CUTANA CUT&RUN and library prep kits from Epicypher (14-1048 and 14-1001), with slight modifications. BMDMs were fixed with 0.1% formaldehyde (ThermoFisher, 28908) for 1 minute, then quenched with glycine (CST, 7005S). Nuclei were isolated with CUTANA Nuclei Extraction Buffer (Epicypher, 21-1026). Nuclei were incubated with SPOP (Genentech, clone F5) or H3K27me3 (CST, 9733S) antibodies overnight. DNA was de-crosslinked by addition of 0.1% SDS and 0.23 μg/μL proteinase K (CST, 10012S), and incubation overnight at 55°C. Libraries were pooled and sequenced on NovaSeq 6000 (Illumina) to generate 20 million paired-end 50-base pair reads for each sample. FASTQs were aligned to the mouse reference genome (GRCm38/mm10) using the Bowtie aligner (version 2.2.6) ^61^. Picard (version 1.126) was used to mark duplicate reads. Peaks were called using MACS2 (version 2.1.0) ^52^ and were filtered to remove blacklisted regions. DiffBind (version 3.16) ^53^ R package was used to perform differential peak analysis relative to *Spop*^-/-^ samples. ChIPSeeker ^56^ was used to annotate peaks. SoGGI ^57^ and Profileplyr ^58^ were used to generate heatmaps and K-means clustering. WashU epigenome browser ^60^ was used to plot the locus specific CUT&RUN read enrichment.

## Acknowledgments

We thank members of the Dixit laboratory and Manolis Pasparakis for advice and discussions; Mariela del Rio Pantle and Fermin Gallardo-Chang for animal husbandry; our colleagues in the Genetically Engineered Animals Program for allele design, creation and screening; our colleagues in Animal Resources for technical assistance; Jessica Mills, Alison Huynh, Victor Nunez, Weibin Liang, Yuxin Liang, and the histology and sequencing laboratories for technical assistance.

## Author contributions

CC and EM designed and performed experiments. RR analysed sequencing data. JJ, KH, and JDW developed and analysed histological data. VK and BD performed SMARCA4 ChIP-seq assays. MS, CC, and SD performed scRNA sequencing. RN and RK developed SPOP antibodies. KA, ZT, and ID performed protein purification. KN, NK and VMD contributed to experimental design. CC and KN wrote the paper with input from all authors.

## Author information

Competing interests statement: all authors were employees of Genentech. Correspondence and requests for materials should be addressed to.

